# Drug-like antibody design against challenging targets with atomic precision

**DOI:** 10.1101/2025.11.29.691346

**Authors:** Chai Discovery Team, Jacques Boitreaud, Robert Chen, Jack Dent, Lucy Fairweather, Danny Geisz, Matthew Greenig, Nate Boyd, Jinay Jain, Brady Johnston, Matthew McPartlon, Joshua Meier, Neil Patil, Zhuoran Qiao, Alex Rogozhnikov, Nathan Rollins, Nikitha Vicas, Paul Wollenhaupt, Kevin Wu, Andy Yeung

## Abstract

Computational antibody design has seen rapid progress, with high success rates enabling direct translation to characterization without any high-throughput screening required. In this work, we markedly expand the scope of *de novo* antibody design by applying our state-of-the-art Chai-2 platform to design drug-like antibodies in full-length monoclonal format. We find that >86% of these full-length mAbs have strong developability profiles on par with therapeutic antibodies. We further show that experimentally determined structures of Chai-2 designs closely match their *in silico* predictions, demonstrating that Chai-2 produces atomically accurate models of designed antibodies. Building on these foundational capabilities, we showcase two potential applications of Chai-2 against different targets: designing functional antibodies mediating GPCR agonism, and highly specific antibodies selectively binding tumor-specific neoepitopes. Taken together, this work brings new flexibility to modern discovery pipelines, accelerating the path from *in silico* design to functional validation across both conventional and challenging targets. Beyond reducing the cost and timelines associated with large screening campaigns, *in silico* design can now open new frontiers for creative, targeted therapeutics that address unmet clinical needs.

## 1 Introduction

In the forty years since the clinical approval of the first monoclonal antibody [1], antibodies have become one of the most effective therapeutic modalities, constituting nearly a third of all newly approved molecules by the FDA [2] owing to their favorable properties including high specificity, excellent pharmacokinetics, and bioavailability. Despite their success, traditional screening-based antibody discovery remains resource-intensive and time-consuming, and often requires extensive downstream optimization after an initial binder is found.

The past year has seen remarkable advances in computational methods for *de novo* protein design, particularly for antibodies [3, 4, 5, 6, 7], demonstrating controllable *in silico* design of antibodies with high binding success rates. However, these works still have several limitations. First, their shared focus on designing single domain antibodies rather than the full length immunoglobulin (IgG) format limits their applicability to therapeutic development. Furthermore, while achieving target binding is essential, true drug candidates must also satisfy multiple developability constraints. Additionally, while several existing models can be prompted with epitopes, they have not yet demonstrated that their designs match reality at atomic scale. This point is underscored by [3] who show that a designed VHH met its biochemical design criteria, yet cryo-EM revealed a markedly different binding orientation than their model predicted.

In June 2025, we released Chai-2 [7], the first model to design antibodies with double-digit success rates. Soon after, we also discovered that the model can design drug-like antibodies against challenging targets with atomic precision. The goal of this whitepaper is to share those results, thereby bridging the gap between proof-of-concept *in silico* binder design and therapeutic viability. Specifically, we demonstrate that Chai-2 can design antibodies as full-length IgGs while maintaining high hit rates established previously. These designed antibodies exhibit developability properties comparable to clinically used therapeutics. Using cryo-EM, we further show that Chai-2’s predicted binding orientations are recapitulated with high accuracy, providing atomic-level confidence in the modeled interactions from the very first hit.

Beyond developability and structural validation, we show that Chai-2 can successfully tackle biologically important and historically challenging target classes, expanding the scope of computational antibody design into areas previously considered inaccessible. One such class of biological systems is G protein-coupled receptors (GPCRs), a large group of multipass membrane proteins whose conformational flexibility, challenging expression, low surface exposure, and limited structural characterization have made them notoriously difficult targets for antibody discovery and rational design. We used Chai-2 to design antibodies against six GPCRs. Not only did the model generate validated binders to all six targets, but it also designed functional agonists to two distinct GPCRs. To our knowledge, these represent the first cases of functional GPCR agonists designed *in silico*, without the use of library-based screening. We also applied Chai-2 to the challenging task of targeting tumor-specific neoepitopes presented by peptide–MHC (pMHC) complexes. We find that Chai-2 is able to design antibodies that discriminate single-residue mutations within the cancer peptide and avoiding binding to wild-type counterparts. These results demonstrate the model’s fine-grained controllability and its precise, atomic-scale understanding of binding interactions.

To summarize our main contributions: Chai-2 can design full-length IgGs with therapeutic-grade developability and precise, atomically steerable binding modes, establishing a viable platform for true end-to-end antibody development. Our case studies featuring functional agonist design for GPCRs as well as single-residue-specificity binders for peptide-MHCs are a glimpse of what is possible when drug design becomes programmable.

## 2 Results

### 2.1 Zero-shot design of full-length mAbs

We previously demonstrated that Chai-2 can design VHHs and scFvs with high success rates. We tested whether the scFv hits could be readily reformatted into full-length mAbs, which are typically used in therapeutic development. Nearly every scFv hit (93%) retained activity in this format, indicating that the original VH-VL sequence design transfers effectively between formats. Altogether, reformatting yielded 88 IgGs, which we next advanced for developability assessment.

### 2.2 Designs exhibit developability comparable to clinical therapeutics

To succeed as therapeutics, antibodies must not only bind their target with high affinity, but also meet key developability criteria: the physicochemical properties that enable production, storage, and administration of the molecule at scale. Poor developability profiles can lead to low manufacturing yields, short shelf-life, and impaired bioactivity due to aggregation or immunogenicity, rendering an otherwise functional drug significantly less useful in the clinical setting. Recent literature has identified high-throughput biophysical assays that serve as early indicators of such issues during hit screening [8, 9, 10, 11, 12]. These assays measure key properties that are predictive of drug viability, including yield and product quality (SDS-PAGE, SEC-HPLC, DLS), thermodynamic stability (Tm, accelerated-stability SEC-HPLC), hydrophobicity (HIC, SGAC-SINS, SMAC), aggregation propensity (AC-SINS, CIC, CSI), and polyspecificity (BVP ELISA, PSR, multi-antigen ELISA).

To assess Chai-2’s potential to design functional antibody therapeutics, we sought to characterize the developability properties of the model’s designs across a panel of industry-relevant biophysical assays. To benchmark against existing molecules, we also screened 40 clinical-stage biosimilar controls, comprising 20 IgGs and 20 VHHs. IgG controls were selected among those in the Jain et al. [8] dataset and available in the Sino Biological reagent catalog to span the range of developability profiles observed in clinical therapeutics. VHHs controls were selected from clinical-stage nanobodies in Thera-SAbDab ([13]). IgGs were expressed as biosimilars, and VHHs were formatted as human IgG1 Fc fusions (VHH-Fcs). All antibodies were produced in HEK293 cells at a 100-mL scale and purified by Protein A column with no further polishing. Samples with adequate monomer purity (≥90% by size-exclusion chromatography (SEC)) were advanced to biophysical assays: thermal stability (NanoDSF), polyreactivity (BVP-ELISA), polydispersity (DLS), hydrophobicity (HIC-HPLC), and self-association (AC-SINS). In total, 19/20 IgG biosimilar controls, 81/88 Chai IgGs, 17/20 VHH-Fc controls, and 17/27 Chai VHH-Fcs were advanced. Note, although low purity may reflect intrinsic molecular liabilities, it may also arise from variability in transient transfection; repeat expression would be required to differentiate these possibilities.

We then applied a widely used scheme proposed in [8, 10], defining “green flag” criteria that identify which designs have promising drug-like properties. First, we benchmarked our assay measurements versus published values for the biosimilar controls: NanoDSF Fab *T_m_* showed good agreement (Figure S1A), HIC-HPLC retention times are slightly extended but show tight linear correlation, BVP ELISA was notably more sensitive but allowed for strong precision-recall of flags, and AC-SINS was less sensitivity but again allowed for good precision-recall of flags. We therefore defined flag thresholds based on [8] with small scaling adjustments for our assay sensitivities. These calibrations are detailed in Figure S1A and the final thresholds used for the four assays are summarized in Table 1.

**Table 1.** Biophysical assay green flag definitions.

| Assay | Green Flag threshold | References |
| --- | --- | --- |
| NanoDSF | $> 64.2^\circ\text{C}$ Fab $T_m$ | Jain et al. [10] |
| | $> 60^\circ\text{C}$ VHH $T_m$ | Valdés-Tresanco et al. [14] |
| HIC-HPLC | $< 12.21$ min retention time | Jain et al. [10] |
| BVP ELISA | $< 29.8$ fold-over-background | Jain et al. [10] |
| AC-SINS | $< 3$ nm shift | Jain et al. [10] |

This analysis showed that a majority of Chai-2 designs achieve three or more “green flags”: biophysical measurement meeting or exceeding widely-accepted thresholds for favorable developability in clinical-stage antibodies [8] (Figure 2A). Figure 2B displays the distribution of these individual biophysical properties, displayed alongside the flag thresholds defined for each assay. Across measurements, Trastuzumab (star) serves as an example of an antibody with a strong developability profile, whereas Ixekizumab (triangle), which is flagged in three assays, represents a poorly-behaved molecule.

**Figure 1.**
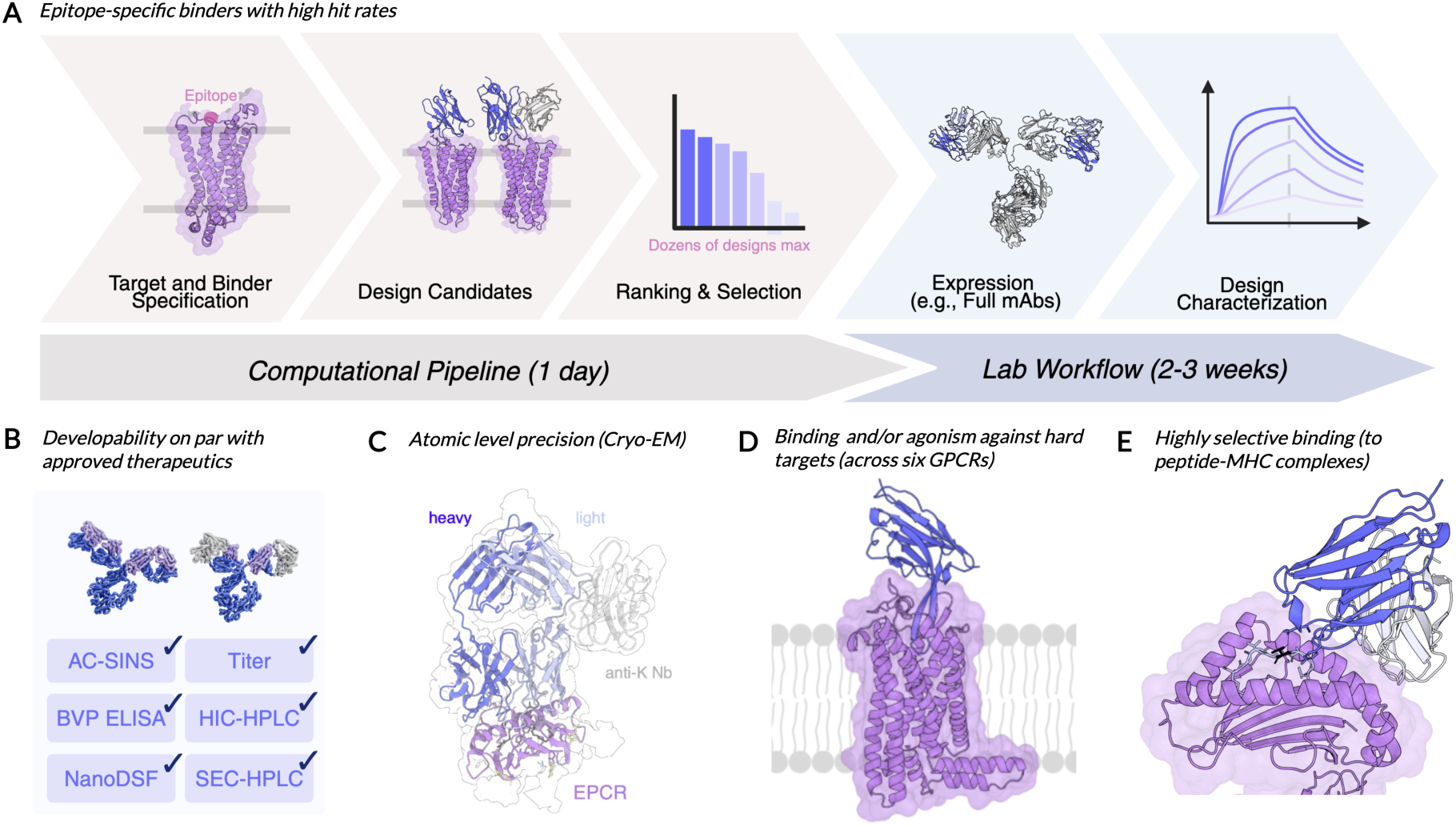
Overview of *in silico* design and experimental validation. (A) Given a target and epitope, Chai-2 designs antibodies and proposes a focused set of candidates. These are then expressed with a variety of formats and experimentally characterized. Beyond binding rates, we characterize developability (B) and validate that experimentally resolved structures match Chai-2’s designs (C). We also leverage Chai-2 to design state-of-the-art antibodies for GPCRs (D) and peptide-MHC complexes (E).

**Figure 2.**
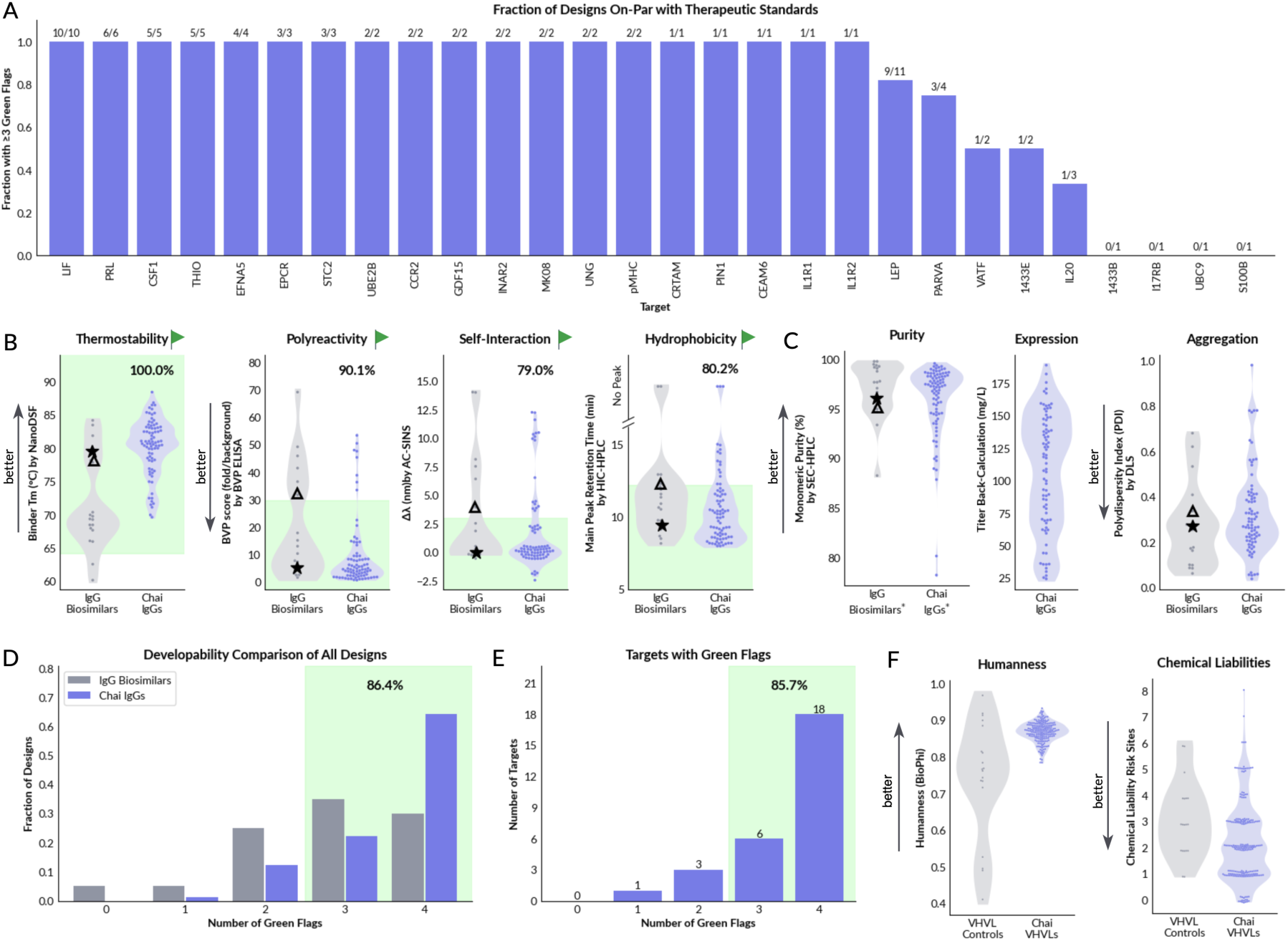
Chai-2 designs exhibit strong developability. All results are experimentally determined in the wet-lab, except for panel F. (A) Fraction of designs that have 3 or more developability green flags by target, annotated with absolute fraction. For 24/28 targets, we recover antibodies with >3 green flags. (B) Distributions of core developability properties of Chai IgGs and biosimilars with therapeutic standard thresholds annotated. Above each violin plot, we report the percentage of designs that pass the corresponding threshold. In each case, over 79% of designs pass each filter. (C) Additional developability lab measurements. Asterisks indicate the purity plot shows all samples; only samples >90% purity advanced to the other assays plotted. (D) Number of green flags as a fraction of antibodies. (E) Number of green flags for the best-scoring design per target. (F) Humanness and chemical liability metrics, computed *in silico*. Trastuzumab (star) serves as an example of an antibody with a strong developability profile, whereas Ixekizumab (triangle), which is flagged in three assays, represents a poorly-behaved molecule.

Chai-designed IgGs exhibit excellent thermostability, with 100% of designs exceeding the 64°C cutoff. They also demonstrate highly favorable polyreactivity, self-association, and hydrophobicity profiles, with 90%, 79%, and 80% of designs meeting the respective thresholds. Notably, Chai IgGs exhibited melting temperatures markedly exceeding those of the biosimilar control set. This trend in part reflects our deliberate selection of the well-behaved VH3-23 and VH3-66 frameworks, which are associated with high thermostability in the Jain et al. [8] dataset (Figure S1B). Yet even compared to framework-matched controls, Chai-2 designs prove to have above-average melting temperatures. This potentially suggests an emergent property of *de novo* designed antibodies, mirroring the high stability characteristic of *de novo* and inverse folding -derived sequences.

In addition to the flag analysis, several more properties were characterized and shown in Figure 2C: back-calculated titer (yielded mass after ProA purification divided by expression volume), monomer purity by analytical SEC, and polydispersity by DLS. Although transient expression results in generally high variation in titer, all designs show high double-digit mg/L titer or better, suggesting that the designs allow for performant expression. Note, differences in culture duration prevent direct comparison of titers between the biosimilars and the designs (see Methods for details). The SEC and DLS measurements show that Chai-2 designed IgGs express at high purity with low aggregation, comparable to the clinical IgG controls.

Figure 2D shows that more than 85% of Chai-2 IgG designs have three or more green flags, with the majority of designs also passing all four checks. Likewise, Figure 2E shows that for 24 of 28 antigens targeted by these 81 IgG molecules, we found at least one corresponding design with at least three green flags. The remaining four targets each had only one design tested; hence, additional screening is likely to further improve the per-target success rate. Similarly, Chai-designed VHHs exhibit developability profiles comparable to clinically developed VHHs (Figures S2 and S3). Importantly, Chai-designed IgGs also exhibit a high degree of humanness and show a low number of predicted chemical liabilities (Figure 2F).

These results demonstrate that Chai-2 can reliably generate functional antibody candidates with favorable developability profiles directly from computational design. Historically, therapeutic antibody development has required extensive experimental screening and iterative engineering cycles to simultaneously optimize binding and biophysical properties [8]. However, the combination of high stability, low aggregation propensity, high purity, strong humanness scores, and minimal predicted chemical liabilities indicates that the model’s designs already serve as viable starting points for preclinical development, without requiring further experimental iterations. By generating designs on the first iteration that meet standard developability criteria, Chai-2 substantially compresses development timelines while maintaining the quality standards required for clinical progression.

### 2.3 Designed 3D interactions match experimental structures with atomic-scale precision

To validate the structural accuracy of our antibody designs, we applied cryo-EM to solve the experimental 3D structures of 5 positive binders: EPCR_design_17, EFNA5_design_10, CSF1_design_6, S1433B_design_3, and IL20_design_9. Each design was reformatted as a Fab and complexed with its respective antigen. The resulting cryo-EM maps ranged from 2.9 Å to 3.9 Å in resolution (gray volumes in Figure 3 A-E). Atomic coordinates (colored cartoons in Figure 3 A-F) were solved from the density volumes using AlphaFold-2 predicted apo-structures for unbound antigen and unbound antibody Fab framework as refinement starting points. No CDR conformation or binding orientation information was provided during refinement, ensuring that no folding model-derived structural information influenced the fitting of the antibody-antigen interaction.

**Figure 3.**
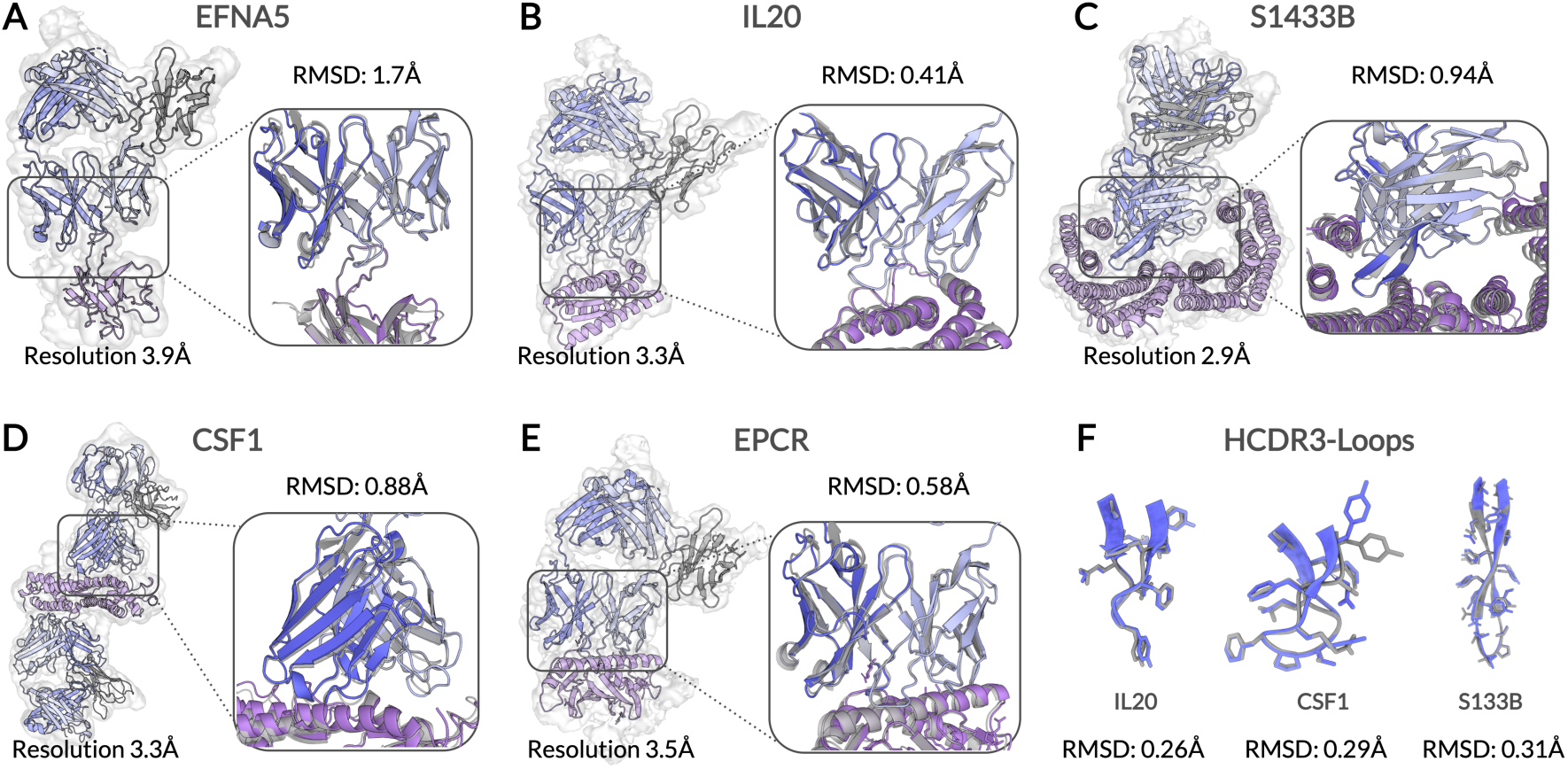
Cryo-EM structures overlaid with ***in silico*** designed structures. Structures are shown for five designs (A-E), with shaded volumes indicating experimentally resolved volumes by cryo-EM, and cartoons indicating computationally predicted and experimentally solved structures for designs. Resolution of each structure is indicated in the corresponding panel, as well as the RMSD error comparing experimental (colored) and computational structures (grayscale) across the antibody-antigen complex (computed for VH-VL and antigen *α*-carbons). (F) shows heavy CDR3 loops for three structures, with gray indicating Chai-2 predicted pose and blue showing experimentally resolved structure.

The all-atom cryo-EM structures exhibited striking agreement with the Chai-2 design models, with global RMSD values ranging from 0.41 Å for S1433B (Figure 3B) to 1.7 Å for EFNA5 (Figures 3 and S5). Detailed analysis (Figure 3F, Figures S4 to S8) reveals that 4 of 5 have sub-angstrom accuracy over the epitope-paratope interface (RMSD from 0.54 Å to 0.71 Å; the outlier, CSF1 is 1.9 Å). Remarkably, HCDR3, the antibody region most unique between designs, shows sub-angstrom accuracy for all 5 designs (HCDR3 RMSD from 0.26 Å to 0.39 Å). Overall, these data show that all five of the antibodies we characterized by CryoEM bound precisely to their intended epitopes and recapitulate nearly all intended interactions, demonstrating that Chai-2 is capable of designing antibodies with near-atomic precision. Extended data showing accuracy of epitopes, paratopes, CDR loops, and 2D cryo-EM class averages can be found in Figures S4 to S8.

### 2.4 Designing antibodies binding and mediating GPCR agonism

#### 2.4.1 Consistent *in silico* discovery of GPCR-binding antibodies

One-third of all approved therapeutics target a G-coupled protein receptor (GPCR), making them one of the most valuable therapeutic targets [15]. However, of the over 500 approved GPCR-targeted agents, only 3 (mogamulizumab, erenumab, and talquetamab) are antibodies [2], none of which exhibit agonistic behavior that directly modulates receptor signaling. A major limitation is the difficulty of generating conformationally active receptor reagents for traditional antibody discovery campaigns.

Chai-2, as a computational method with high success rates, can avoid high-throughput screening and thereby sidestep many experimental challenges associated with GPCRs. Here, we selected GPRC5D (G protein-coupled receptor family C group 5 member D; 9IMA) and a panel of class A chemokine GPCRs [16], namely CCR2 (7XA3), CCR5 (6AKX), CCR8 (8KFX), CXCR4 (8U4R), and CXCR6 (8T34), for model testing. We prompted Chai-2 to target surface epitopes on all GPCRs, except for CCR8, CXCR4 and CXCR6, where we specifically instructed the model to target the orthosteric pockets. Additionally, while most receptors were targeted with single-domain VHHs, we also tested a full-length IgG approach for two receptors: CCR2 and CCR5.

Table 2 summarizes the results of these screens using various assays. Chai-2 successfully generated hits against all five GPCRs tested, achieving a target hit rate of 100% with per-target hit rates ranging from ∼4% to 87% (median: 24%), despite evaluating only double-digit designs per receptor, per format. Furthermore, our full-length IgG designs are similarly successful, with IgG designs to CCR2 exhibiting a similar hit rate as VHH designs, and all IgG designs to CCR5 exhibiting binding. While previous works have shown that computational methods coupled with high throughput screening can design VHHs binding GPCRs [17], Chai-2 takes this a step further by designing full-length IgGs to these challenging targets, all while testing fewer than 50 designs.

**Table 2.** GPCR binding and functional hit rates. Detailed results are included in Figure S9. For GPRC5D, function is achieved by binding and designs were not intended to be agonists. For the other targets, function refers to agonism.

| Target | Format | Screen | Number tested | Number of binding hits | Number of functional hits |
| --- | --- | --- | --- | --- | --- |
| CCR8 | VHH-Fc | SPR | 10 | 5 | 1 |
| CXCR4 | VHH-Fc | ELISA | 19 | 2 | 1 |
| CXCR6 | VHH-Fc | ELISA | 23 | 1 | 0 |
| GPRC5D | VHH-Fc | FCM | 73 | 35 | Not tested |
| CCR2 | VHH-Fc | BLI | 40 | 10 | Not tested |
| CCR2 | IgGs | BLI | 45 | 8 | Not tested |
| CCR5 | IgGs | BLI | 13 | 13 | Not tested |

**Table 3.** Cryo-EM data collection, processing, and refinement statistics for EFNA5/Fab, EPCR/Fab, CSF1/Fab, IL20/Fab, and 1433B/Fab complexes.

|  | EFNA5/Fab<br>complex | EPCR/Fab<br>complex | CSF1/Fab<br>complex | IL20/Fab<br>complex | 1433B/Fab<br>complex |
| --- | --- | --- | --- | --- | --- |
| **Data collection and processing** |  |  |  |  |  |
| Magnification | 130,000 | 130,000 | 105,000 | 130,000 | 130,000 |
| Voltage (keV) | 300 | 300 | 300 | 300 | 300 |
| Electron exposure ($e^-/\text{\AA}^2$) | 50 | 50 | 48 | 48 | 48 |
| Defocus range ($\text{\AA}$) | -1.0 to -2.2 | -1.0 to -2.2 | -1.0 to -2.2 | -1.0 to -2.2 | -1.0 to -2.2 |
| Pixel size ($\text{\AA}$) | 0.97 | 0.932 | 0.824 | 0.932 | 0.932 |
| Symmetry imposed | C1 | C1 | C2 | C1 | C1 |
| Initial particle images | 4,522,190 | 4,272,470 | 2,714,578 | 1,591,991 | 3,468,127 |
| Final particle images | 357,683 | 428,640 | 1,030,990 | 518,312 | 1,246,613 |
| Map resolution |  |  |  |  |  |
| FSC threshold ($\text{\AA}$) | 0.143 | 0.143 | 0.143 | 0.143 | 0.143 |
| Map resolution ($\text{\AA}$) | 3.9 | 3.5 | 3.3 | 3.3 | 2.9 |
| **Refinement** |  |  |  |  |  |
| Initial model used (PDB) | Ab-Initial | Ab-Initial | Ab-Initial | Ab-Initial | Ab-Initial |
| Model resolution |  |  |  |  |  |
| FSC threshold ($\text{\AA}$) | 0.143 | 0.143 | 0.143 | 0.143 | 0.143 |
| Map sharpening $B$-factor ($\text{\AA}^2$) | -167.0 | -174.6 | -191.5 | -145.8 | -110.2 |
| **Model composition** |  |  |  |  |  |
| Non-hydrogen atoms | 5,286 | 5,653 | 9,912 | 5,029 | 7,505 |
| Protein residues | 802 | 841 | 1,392 | 707 | 1,017 |
| Ligands | 0 | 1 | 0 | 0 | 0 |
| $B$-factors ($\text{\AA}^2$) | 1.72 | 0.50 | 0.50 | 2.57 | 7.89 |
| **R.m.s. deviations** |  |  |  |  |  |
| Bond lengths ($\text{\AA}$) | 0.006 | 0.005 | 0.006 | 0.006 | 0.006 |
| Bond angles ($^\circ$) | 0.940 | 0.940 | 0.920 | 0.910 | 0.910 |
| **Validation** |  |  |  |  |  |
| MolProbity score | 1.70 | 1.69 | 1.31 | 1.23 | 1.20 |
| Clashscore | 8.10 | 8.17 | 5.68 | 4.56 | 4.13 |
| Rotamer outliers (%) | 0.00 | 0.00 | 0.00 | 0.00 | 0.00 |
| **Ramachandran plot** |  |  |  |  |  |
| Favored (%) | 96.13 | 96.32 | 98.04 | 98.43 | 98.11 |
| Allowed (%) | 3.87 | 3.68 | 1.96 | 1.57 | 1.89 |
| Outliers (%) | 0.00 | 0.00 | 0.00 | 0.00 | 0.00 |

#### 2.4.2 Discovery of cross-reactive binders to an orpan GPCR

GPRC5D is an orphan GPCR that has been identified as a promising therapeutic target in multiple myeloma [18, 19]. Notably, the recently solved GPRC5D structures were not included in Chai-2’s training dataset, providing a natural test case for evaluating model performance on an unseen target. In this setting, Chai-2 achieves a surprisingly strong 48% hit rate (35 hits / 73 tested designs, Table 2 and Figure S9). Structural clustering of all designs revealed convergence on a common epitope with highly similar binding orientations to talquetamab. We also confirmed cell-surface binding of these designs by FACS (Figure 4B) and nanodiscr-constituted human and cynomolgus GPRC5D. Sixteen of the twenty-four tested showed clear SPR binding to both orthologs, with the top hit GPRC5D_design_47 exhibiting *K_D_*values of 189 nM to human and 334nM to cyno nanodisc material (Figure 4C). Collectively, our findings reveal that Chai-2 can reliably generate epitope-specific antibodies against a challenging class C GPCR target, underscoring the model’s ability to design for loop-dominant epitopes and establishing a strategy for targeting orphan receptors of therapeutic relevance.

**Figure 4.**
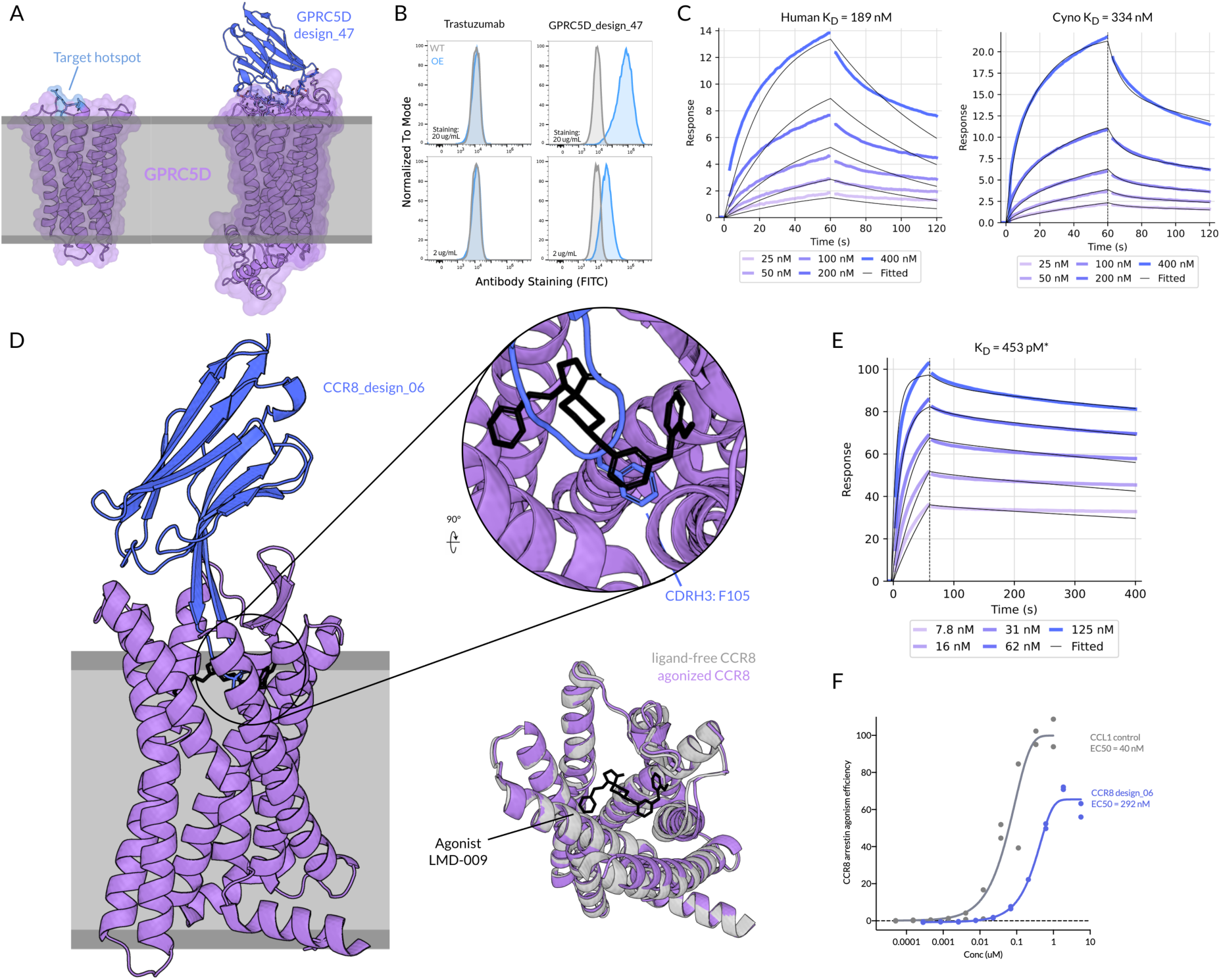
Binding and functional hits to GPCRs. (A) Structure of GPCR5D (PDB: 9IMA) with target hotspot (left) and predicted structure of VHH GPRC5D_design_47 (right) identified by testing desings with flow cytometry. The blue-highlighted hotspot is predicted to be cyno cross reactive. (B) Flow cytometry binding of GPRC5D_design_47 to GPRC5D overexpression and wild-type CHO-K1 cells. Healthy singlet cells were first gated prior to plotting FITC-A. A shift was observed, demonstrating potent binding. (C) Monovalent affinities of GPRC5D_design_47 to nanodic-solubilized human and cynomolgus monkey GPRC5D proteins were determined to be 189 and 334 nM respectively. (D) Chai-2 predicted interaction of VHH CCR8_design_06 to LMD-009-agonized CCR8 (PDB: 8KFX). CCR8 without guanine nucleotide-binding protein (G_i_) is depicted in purple, LMD-009 is depicted in black, and the designed VHH is depicted in blue (left). In the inset, the F105 side chain in CCR8_design_06’s CDRH3 is shown. A top view of ligand-free CCR8 (PDB: 8KFZ) and LMD-009-agonized CCR8 are depicted (bottom right). (E) Bivalent affinity of CCR8_design_06 to CCR8 nanodisc-solubilized protein was determined to be 453 pM via SPR. (F) PathHunter CCR8 arrestin agonism assay was performed with CCR8 designs and CCL1 control ligand. The dose response curve of CCR8_design_06 compared to CCL1 is plotted, to which a 4-parameter nonlinear fit was performed to determine the EC_50_ values. EC_50_ values of 292 nM and 40 nM were respectively determined. *K_D_*values marked with * are measured with avidity.

#### 2.4.3 Discovery of GPCR agonists to chemokine receptors CCR8 and CXCR4

Functional activation of GPCR signalling – also known as agonism – has emerged as a successful therapeutic strategy across a wide range of drugs, including GLP-1 agonists for diabetes and obesity, *β*2-adrenergic receptor agonists for asthma, and PTH receptor agonists for osteoporosis [20]. Small molecule and peptide agonists typically achieve this by targeting the orthosteric site, a concave epitope buried inside the transmembrane bundle [21], but traditional antibody discovery methods typically struggle with such deep, concave pockets. Therefore, we applied Chai-2 to design functional VHH agonists specifically targeting the buried orthosteric site in three therapeutically relevant GPCRs (CCR8, CXCR4, and CXCR6).

For CCR8, a chemotaxis mediator implicated in cancer immunology [22, 23], we assessed all 10 designs along with positive control native ligand CCL1 in a *β*-arrestin recruitment assay using the PathHunter platform. Our results show that two designs exhibited detectable activity, with one construct, CCR8_design_06, showing clear agonism with an EC_50_ of 292 nM and a modestly reduced E_max_ relative to the CCL1 control (Figure 4F), within an order of magnitude of the positive control EC_50_ of ∼40 nM. Hence, from only ten initial designs, we identified a bona fide agonist antibody that functionally engages and activates the receptor.

Structural modeling of CCR8_design_06 in complex with CCR8 (Figure 4D) suggests that CCR8_design_06 mimics the mechanism of action of small-molecule agonist LMD-009 described in the active-state CCR8-G_i_ cryo-EM structure [24] by positioning an CDRH3 aromatic residue, F105, around the conserved YYE triad. In fact, several other designs were also predicted to similarly insert an aromatic side chain into this pocket, but CCR8_design_06 stood out kinetically: it exhibited the slowest dissociation rate (*k*_off_ ≈4.06 × 10*^−4^s^−1^*) among the panel (Figure 4E). These observations suggest that sustained receptor engagement, rather than pocket binding alone, may be critical for driving productive CCR8 activation in this design.

We next evaluated designs against CXCR4, a receptor that signals through CXCL12 to regulate chemotaxis, stem-cell homing, and immune-cell trafficking, with functional implications for key processes such cancer metastasis, immune regulation, and HIV infection [25]. 19 designs were tested with a G_i_-coupled cAMP cell-based functional assay, which measures CXCR4-mediated suppression of cAMP as a hallmark of canonical signaling. We observed the highest-affinity binder (CXCR4_design_06, Figure S9D) exhibiting partial agonist activity in the cAMP assay, with an estimated EC_50_ of ∼164 nM compared to 0.83 nM for CXCL12 (Figure Figure S9E). Compounding on our functional designs for CCR8, these results provide additional proof-of-concept that Chai-2 can design antibodies that elicit measurable agonism in chemokine receptors.

### 2.5 Selective targeting of peptide-MHC complexes

Next, we turned to another challenging target class: peptide-Major Histocompatibility Complexes (pMHC). pMHCs comprise short peptides derived from intracellular proteins that are loaded onto MHC molecules and displayed on the cell surface. This process allows the immune system to recognize intracellular proteins, which is especially relevant when peptides from disease-specific or mutated drivers, such as oncogenic mutations or pathogen-derived proteins, become accessible once presented extracellularly [26]. Antibodies or engineered binders specific to these pMHCs can therefore engage intracellular disease drivers that would otherwise be inaccessible, offering new therapeutic opportunities in oncology and infectious disease.

However, generating high-quality pMHC binders remains challenging. Due to the highly conserved nature of MHC molecules, the specificity of a particular pMHC is largely encoded by only two to four distinct exposed peptide residues, making discrimination among closely related pMHCs particularly difficult. Compounding this, the MHC-I groove forms a concave surface that is not well-suited to conventional antibody paratopes [27], limiting their ability to engage all key peptide residues simultaneously. This, in turn, makes high-affinity, selective binding difficult to achieve. Computational design faces additional hurdles, including accurately modeling peptide dynamics and conformational variability to capture true pMHC structure during binder generation.

In this study, we tested a limited number of Chai-2 designs against three therapeutically relevant peptide-MHC complexes: HLA-A*02:01-p53 R175H (HMTEVVRHC), HLA-A*02:01-gp100 (YLEPGPVTV), and HLA-A*03:01-KRAS G12V (VVVGAVGVGK, Figure 5A, left panel), testing 27, 48, and 50 designs respectively. Only the A*03:01-KRAS G12V campaign produced hits, yielding two binders, whereas the p53 R175H and gp100 screens produced no detectable binders (Figure S10). Part of the failure likely reflects assay limitations, as the p53 R175H positive control (AB330) generated only marginal signal in our ELISA format (Figure S10), indicating a fundamental limitation of available assay reagents and sensitivity in this context.

**Figure 5.**
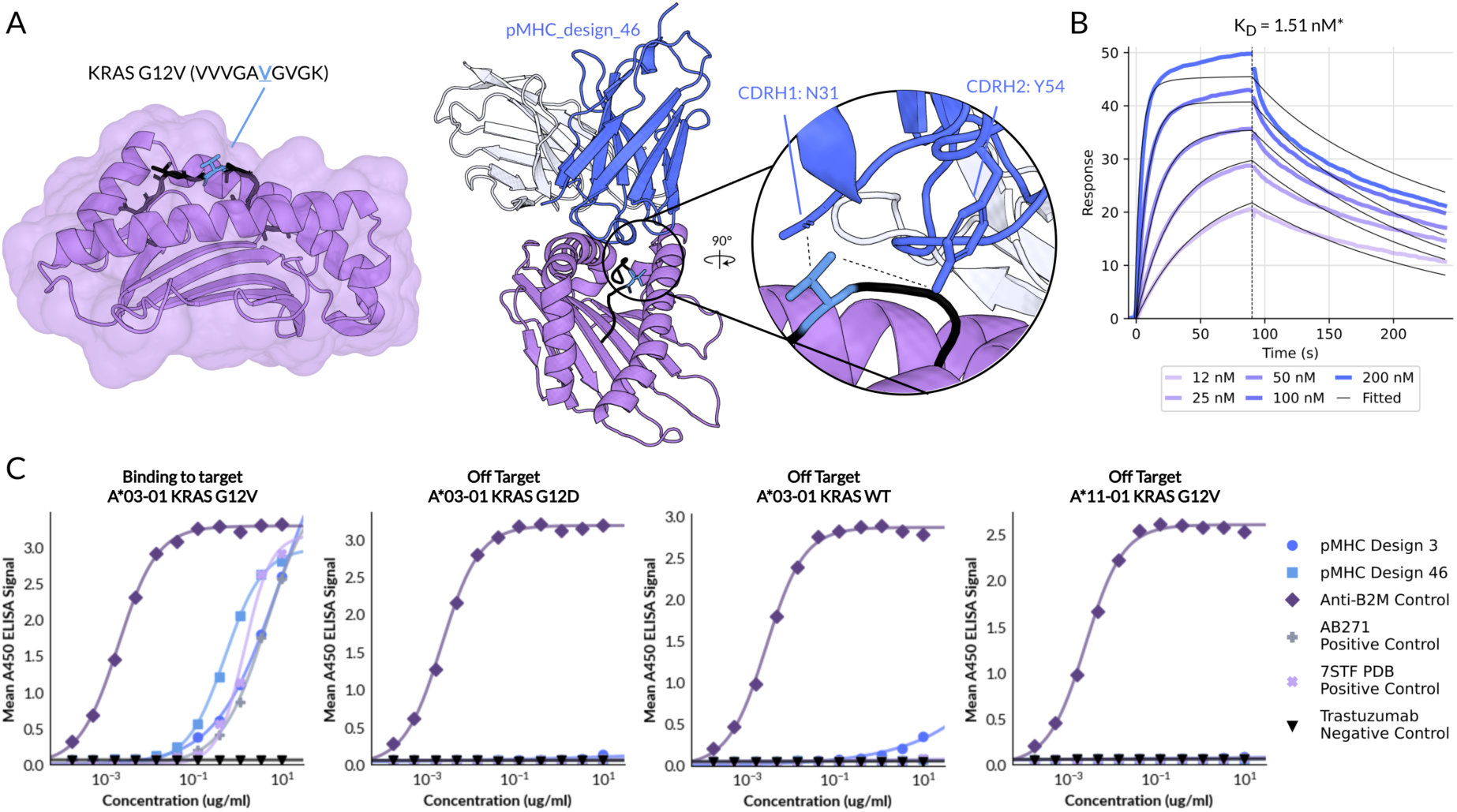
Structure modeling and characterization of pMHC-binding antibody. (A) Structure of HLA-A*03:01 bound to KRAS G12V (VVVGAVGVGK) (PDB: 7STF), with B2M cropped for visualization (left). Depicted design_46 (right) was discovered via ELISA screening (Figure S10C) and was characterized in further assays. In the inset, the G12V, N31 CDRH1 and Y54 CDRH2 side chains from design_46 are depicted. The dashed lines indicate measured distances of 4.2 and 4.8Å between N31 and Y54 respectively to the valine C3’ and C4’ carbons. (B) Bivalent affinity of design_46 to His-immobilized HLA-A*03:01 & B2M & KRAS G12V (VVVGAVGVGK) was determined to be 1.51 nM via SPR. (C) ELISA binding was performed for design_3 and design_46 across four peptide MHC reagents: design target HLA-A*03:01 & B2M & KRAS G12V (VVVGAVGVGK) and off-targets HLA-A*03:01 & B2M & KRAS G12D (VVVGADGVGK), HLA-A*03:01 & B2M & KRAS WT (VVVGAGGVGK), and HLA-A*11-01 & B2M & KRAS G12V (VVVGAVGVGK). EC_50_s were calculated from a 4-parameter nonlinear fit and determined to be 60.43 nM, 3.56 nM, 13.8 pM, 30.51 nM, and 9.75 nM respectively for anti-A*03:01 KRAS G12V design_03, design_46, anti-B2M positive control, AB271 positive control, and the positive control binder from 7STF. An EC_50_ for Trastuzumab could not be determined. *K_D_*values marked with * are measured with avidity.

The two KRAS G12V designs (design_46 and design_03) showed dose-dependent binding comparable to positive-control antibodies (Figure 5C). As shown in Figure 5B, design_46 bound recombinant HLA-A03:01-KRAS G12V with an apparent affinity of 1.5 nM. Both designs also displayed strict mutation specificity (Figure 5C), showing negligible binding to wildtype KRAS or KRAS G12D pMHCs, each differing by only a single amino acid substitution; notably, design_46 showed no detectable cross-reactivity even at 10 *µ*g/mL. Binding was additionally allele-specific, as neither antibody recognized HLA-A11:01-KRAS G12V (Figure 5C), confirming that recognition depends on the structural features and peptide presentation of HLA-A*03:01 rather than on the peptide alone.

Structural modeling using Chai-2 further showed that KRAS-G12V_design_46 engages the pMHC in a binding (Figure 5A, right panel) distinct from the known antibody structure in PDB: 7STF (Figure S11), a structure that was not part of Chai’s training data. The model also predicts that two residues, N31 (CDRH1) and Y54 (CDRH2), form key contacts with the valine mutation that serves as the specificity handle, providing a molecular explanation for discrimination against WT and G12D variants.

Together, these results demonstrate that Chai-generated pMHC antibodies can achieve the high degree of allele- and mutation-specific precision required for pMHC therapeutics, with binding profiles that cleanly distinguish among single-residue peptide variants.

### 2.6 Future work and Limitations

In this work, we aimed to further characterize designs from the Chai-2 release in June 2025. To assess the drug-likeness of our molecules, we performed a battery of developability assays, finding that molecules achieve many of the properties required for further advancement. However, future work will require additional in vivo testing to rigorously evaluate the pharmacokinetic and pharmacodynamic properties of the designs.

It should be noted that GPCR and pMHC were selected as antigens for proof-of-concept benchmarks to demonstrate Chai-2’s capabilities on canonically difficult target classes. While we successfully identified binders and observed initial functional activity, these antibodies were designed to validate the model rather than to meet a patient-oriented target product profile (TPP), which would require further optimization, comprehensive developability assessment, more extensive functional characterization (including in vivo testing), and lead refinement. Therefore, while our experiments establish Chai-2 as a frontier generative model for therapeutic antibody design against challenging targets, we also highlight the substantial work that remains necessary to advance designs towards clinical development.

Looking ahead, the reliability of models will only continue to increase. Eventually, models may even generate full development candidates, zero-shot, to a large fraction of targets. The rapid downstream testing afforded by generative models could enable seamless extension into advanced biologic formats such as bispecifics, multispecifics, and antibody-fusion or conjugate modalities, which represent a rapidly expanding share of today’s therapeutic pipeline.

## 3 Discussion

Five months ago, Chai-2 was released as the first method capable of designing antibodies with high success rates in the lab [7]. Here we further validate that Chai-2 can be applied to design atomically-accurate antibodies from scratch in therapeutically relevant mAb formats without high-throughput screening, including for “hard to drug” targets. In addition to achieving high target-binding success rates from testing only dozens of designs, Chai-2 generates antibodies that satisfy multiple properties essential for therapeutic candidates (“developability properties”). Antibodies need to achieve strong developability properties to advance from discovery to the clinic, as molecules must express well, remain stable, exhibit predictable pharmacokinetics and safety, and avoid liabilities such as aggregation or polyreactivity to become viable drugs. Strikingly, our experiments reveal that Chai-2 designs exhibit remarkably favorable profiles, with high thermal stability, low polyreactivity, low hydrophobic binding tendency, and minimal self-interaction, which are superior to those typically reported for previous AI-based approaches and often even exceed those of biosimilar therapeutic antibodies. This highlights the strong potential for Chai-2 antibodies to translate directly into therapeutic development.

It is encouraging that AI-based models can generate high-affinity and developable antibodies against difficult targets in an epitope-specific manner, but it is critical to validate that these binders engage their targets in the intended binding mode in order to directly evaluate the accuracy of the underlying models. RFAntibody [3] underscored this point by showing that a designed VHH against SARS-CoV-2 RBD met its biochemical design criteria, yet cryo-EM revealed a markedly different binding orientation than the design. In contrast, in our work we solved five cryo-EM structures of Chai-2 designed antibodies and found that all five bound their targets with near-atomic agreement to the design models, with RMSDs ranging from 0.41 Å to 1.7 Å. This level of structural concordance demonstrates that Chai-2 can reliably design antibodies with high confidence in the predicted binding mode prior to experimental testing, enabling epitope-specific antibody design. Furthermore, given that our designs are paired with high-confidence structural models, downstream iterative optimization approaches such as affinity maturation and enhancement of biophysical properties can be readily enabled.

In this work, we also tested Chai-2 on challenging target classes, GPCRs and pMHCs, which traditionally require iterative and extensive experimental campaigns for binder discovery. Our results demonstrate that Chai-2 can produce both VHH and full IgG binders across multiple GPCR classes and can even achieve chemokine receptor agonism. This suggests that our model enables the generation of binders to both loop and orthosteric regions, underscoring the adaptability of the design strategy. The identification of a CCR8 agonist from only 10 tested clones provides clear evidence that GPCR agonism is achievable by design. In principle, this extreme precision enables testing directly on relevant cells, including primary cells and tissues from patients, which may improve translational confidence by assessing highly conformation-sensitive receptors in their most native context.

In addition to GPCRs, highly specific pMHC binders hold substantial therapeutic value, enabling precise targeting of intracellular antigens and expanding the range of disease-associated proteins that can be modulated with biologics. Recently, Bennett et al. [3] reported a pMHC binder against the PHOX2B-HLA-C*07:02 complex with micromolar affinity using RFantibody, which tests millions of sequences in a yeast-display library with combinatorial assembly. In contrast, our approach directly designs full-length IgGs with low-nanomolar affinity and recovers mutation and allele-specific binders from a compact set of just 50 designs.

These results suggest that antibody engineering is beginning to shift from laborious empirical screening toward more intentional molecular design. With Chai-2, it is possible to design full-length mAbs with therapeutic attributes and function, before a single experiment is ever run in the wet-lab. This programmability heralds a new era of therapeutics design and could open entirely new frontiers for unmet clinical challenges.

## 5 Methods

### 5.1 Antibody expression and purification

VH and VL gene fragments were ordered and cloned into standard human CH1 IgG1 Fc heavy chain vectors or human CL1 vectors. VHH gene fragments were cloned with a standard junction for format into VHH human IgG1 Fc fusions. All design mAbs were expressed as IgG1. Heterologous expression was performed in HEK293 cells, with transient transfection performed with 1:1 heavy:light transfections or just the VHH-Fc single chain utilizing a PEI-based reagent. After four days of expression, supernatant was clarified. Antibody was purified via pre-packed gravity 2mL column volume gravity columns packed with custom-developed Protein A resin (equivalent to Cytiva 17549803). Supernatant was bound for 90 minutes. Resin was washed three times with 50mM Tris, 100 mM NaCl, pH 8.0. Antibody was eluted with 100mM Glycine, 10mM NaCl, pH 3.0 and promptly neutralized with 1/10th the elution volume of 1 M Tris-HCl, pH 8.5. Antibody was concentrated and buffer exchanged into antibody storage buffer, 1x PBS pH 7.4, and held at 4°C for further use. Purified antibody was assessed for purity on SDS-PAGE and percent monomer was quantified via SEC-HPLC.

The IgG biosimilars were adalimumab, atezolizumab, basiliximab, belimumab, bevacizumab, bococizumab, briakinumab, cetuximab, elgemtumab, elotuzumab, infliximab, ipilmumab, ixekizumab, lintuzumab, ocrelizumab, palivizumab, pembrolizumab, rituximab, trastuzumab, and ustekinumab. The 20 VHH benchmark binders were brivekimig1_VHH1, brivekimig2_VHH1, caplacizumab_VHH1, envafolimab_VHH1, erfonrilimab_VHH2, gacovetug_VHH1, gefurulimab_VHH1, gocatamig2_VHH1, letolizumab_VHH1, lofacimig_VHH1, ozoralizumab_-VHH1, ozoralizumab_VHH2, podentamig1_VHH2, produvofusp_VHH1, rimteravimab_VHH1, sonelokimab2_-VHH1, and tarperprumig_VHH2 [13].

### 5.2 Biolayer interferometry (BLI)

BLI was performed on a ForteBio Octet RED384. In brief, purified antibodies were generally diluted down to 10 ug/mL in BLI running buffer and captured via ProA biosensor (Sartorius 18-5010) for 300 seconds. Single-point affinity screening was performed with antigen usually at 5 uM concentration. Positive binding via response (nm) was generally utilized as a cutoff. A blank biosensor in BLI running buffer was utilized for per-cycle normalization. Binding response was analyzed in Octet® Data Analysis HT software. For a binder to be considered interacting and passed onto 5-point SPR we utilized a cutoff 0.1 nm above the negative control biosensors and greater than 300% over background. PBST (10 mM Na2HPO4, 2 mM KH2PO4, 137 mM NaCl, 2.7 mM KCl, 0.02% Tween-20) with the addition of 0.1% BSA, pH 7.4 was used as the BLI running buffer for all samples.

### 5.3 Surface plasmon resonance (SPR)

SPR was run on a Biacore 8K or T200 depending on instrument availability. For most cases, the antibody was captured via Sensor Chip Protein A (Cytiva 29127555) at a concentration of 5 ug/mL with a 25 second contact time at a flow rate of 10 *µ*L/mL. The target capture level was 200 RU, though with some antibody-antigen pair dependent assay condition tailoring. The antigen concentration sweep was determined from roughly a 10-fold concentration starting point based on the estimated KD from the single-point BLI screen above. Antigen was injected at 30 *µ*L/min for typically 60 seconds followed by a dissociation phase dependent on the antibody-antigen pair, 60-300 seconds. After each cycle, the sensor surface was regenerated via 10 mM glycine-HCl pH 1.5 (Cytiva BR100354). Kinetic profiles were qualitatively assessed to avoid diffusion limited kinetics and exhibit reasonable binding responses in association. 1:1 Langmuir binding curve fits were determined and association, dissociation, and affinity are reported. 1x HBS-EP+ Buffer pH 7.4 (Cytiva BR100826) was used as the running buffer for all samples, and antigens with excess excipient content were exchanged into antibody storage buffer.

### 5.4 Cryo-EM specimen preparation, data collection, and model refinement

Aliquots of antibody-antigen complexes were applied to glow-discharged Quantifoil 300 mesh R 1.2/1.3 Ul-trAuFoil Holey Gold Films. The grids were blotted at 100% humidity and plunge-frozen into liquid ethane using a Vitrobot Mark IV (Thermo Fisher Scientific). Grids were imaged on a 300 kV Titan Krios microscope (Thermo Fisher Scientific) and equipped with a Falcon 4i direct electron detector and a Selectris energy filter. Movies were collected at a magnification of 130,000× and stored in EER format with 1080 raw frames per movie. The dose was set to 50 electrons per Å*^2^*. Automated data collection was carried out using EPU software with a nominal defocus range of –1.0 to –2.2 *µ*m.

All processing was performed in RELION-5.0. The ∼9,555 movies were motion-corrected using the RELION implementation of MotionCor2. The contrast transfer functions (CTFs) of the motion-corrected micrographs were determined using CTFFIND-4.1 and particles were picked using auto-picking. A particle stack selected after 2D classification was used for 3D classification. Subsequent processing involved multiple rounds of 3D classification, 3D auto-refinement, and CTF refinement, yielding reconstructions. Resolution was determined at a criterion of 0.143 Fourier shell correlation (FSC) in the gold-standard refinement procedure. The final map was sharpened using post-processing procedure. Predicted models were docked into the map using UCSF Chimera. Iterative rounds of model building and refinement were performed in CCP-EM-1.4.1 and COOT-0.9.5. Key data collection and refinement parameters are presented below:

### 5.5 Peptide MHC ELISAs

Peptide MHC (pMHC) was coated on 96-well ELISA plates (Jincanhua 1041-007) at 100 uL per well at 2 ug/mL in 1× PBS overnight, sealed at 4°C. Plates were then washed three times and subsequently blocked with 3% BSA in 1× PBS for one hour at room temperature. Plates were then washed five times and primary antibodies were serially diluted and bound to pMHC in 3% BSA in 1x PBST for one hour at 37°C. After this point, plates were then washed three times. Detector antibodies were added: HRP anti-mouse Fc (Jackson 115035003) for anti-B2M antibody (Thermo MA1-19141) or HRP-anti human Fc (Jackson 109035008) for all others, at 1:5000 dilution into 3% BSA in 1x PBST and incubated for one hour at 37°C, oscillating at 600 rpm. Plates were then washed three times. 100 *µ*L of TMB substrate (Biopand TMB-S-004) was then added per well and incubated until sufficient color developed in the B2M plate control. Finally, the reaction was quenched via the addition of 50 *µ*L of 0.5 mol/L sulfuric acid and plates were read on a plate reader at Abs 450. Washes were all performed with 1x PBS with 0.5% Tween-20 (PBST).

### 5.6 Flow Cytometry

Cells were resuspended in 5 mL of pre-cooled staining buffer (PBS + 1% BSA for all steps) and washed twice by centrifugation at 800-1400 rpm × 5min. Cell density was adjusted to 2E7 cells/mL with staining buffer. 5E5 to 1E6 cells per assay were then stained at 100*µ*g/mL of antibody at 30 minutes to 1 hour at 4°C. Cells were washed twice by centrifugation at 800-1400 rpm for 5min. Secondary staining was then performed using FITC anti-human IgG (Sino custom product) at a staining concentration of 20 *µ*g/ml at 30 minutes to 1 hour at 4°C. Finally, cells were washed twice by centrifugation at 800-1400 rpm for 5min with a final suspension to 200 *µ*L of staining buffer prior to analysis on a NovoCyte how cytometer. Trastuzumab (Sino 68046-H002) was used as an isotype control.

### 5.7 Analytical SEC-HPLC

Analytical SEC was performed on an Agilent 1290 with SEC column AdvanceBio SEC 300A 7.8*150mm, 2.7*µ*m pore size. Samples were autosampled and run in a 0.1 M sodium phosphate, pH 6.5 buffer mobile phase with a flow rate of 0.6 mL/min at room temperature. Detection was performed with UV 280 nm. Samples were clarified at 10,000 rpm for 5 minutes, and then supernatant was collected. The column was equilibrated with the mobile phase for 40 minutes, prior to injection of samples. Following program collection, data was analyzed using area normalization methodology to report % purity.

### 5.8 Hydrophobic interaction chromatography HIC-HPLC

HIC-HPLC was performed on an Agilent 1290 with AdvanceBio HIC 4.6*100mm column. 5 *µ*g of IgG samples (at 1 mg/mL) were spiked in with a mobile phase A solution (1.8 M ammonium sulfate and 0.1 M sodium phosphate at pH 6.5) to achieve a final ammonium sulfate concentration of about 1 M before analysis. Then, a linear gradient of mobile phase A and mobile phase B solution (0.1 M sodium phosphate, pH 6.5) was run over 20 min at a flow rate of 1 mL/min with UV absorbance monitoring at 280 nm. Phase C solution (0.1 M sodium phosphate, pH 6.5 with organic agent) was run to clean the column between samples. Following program collection, data was analyzed using area normalization methodology to report HIC retention time for the major peak. If a peak was unresolvable we reported the retention time as an arbitrary 25 minutes [8]. Longer HIC retention times indicate higher hydrophobic interaction tendencies of the test article.

### 5.9 Affinity-capture self-interaction nanoparticle spectroscopy (AC-SINS)

The AC-SINS assay was performed as described previously [28]. Polyclonal goat anti-human IgG Fc capture antibodies and goat non-specific antibodies (non-capture) were buffer exchanged into 20 mM NaAc, pH 4.3, followed by a concentration normalization to 0.4 mg/mL. A 4:1 volume ratio mix of capture:non-capture IgG solution was prepared for 80 capture capacity coating solution. A 9:1 volume ratio was used to mix gold nanoparticle solution with coating solution. After room temperature incubation for 1 h, thiolated PEG (final concentration 0.1 *µ*M) was used to block empty sites on the AuNP. The particle solution was then passed through a 0.22 *µ*m PVDF membrane. PBS at 1/10 of the starting volume was used to elute the particles into the collection tube. 10 *µ*L of the 10X concentrated coated particles were incubated with 100 *µ*L of test antibody solution (50 *µ*g/mL or above in PBS) at room temperature for 1 h in a polypropylene plate, then 100 *µ*L of the resulting solution was transferred into a polystyrene UV transparent plate. Absorbance data was collected on a plate reader from 510 nm to 570 nm in increments of 2 nm. PBS and herceptin were utilized as internal assay controls to assess system and assay run suitability prior to data release. Data is reported as the absorbance shift between captured and non-captured IgG solutions, where self-interacting clones have a larger wavelength shift.

### 5.10 Differential Scanning Fluorimetry (DSF)

NanoDSF was run on a Prometheus NT.Plex. Samples were run from 20-95°C at a ramp rate of 1.5°C/min on automatic fluorescence gain settings. Approximately 10 *µ*L of antibody samples were prepared and added per sample at 0.5 or 1 mg/mL, depending on stock concentration. The first time derivative was analyzed to find the inflection point for *T_m_* utilizing instrument analysis software. The *T_m_* was assigned using the first derivative of the raw data, with the first inflection point *T_m_*1 being reported to match the Jain et al. [8] analysis.

### 5.11 Dynamic Light Scattering (DLS)

DLS was performed on a Zetasizer Pro. After 30 minutes of pre-heating, the assay was set up with Nanoparticle Size and Zeta Potential Analysis options. The sample was mixed and 70*µ*L at 1 mg/mL was transferred to cuvette while avoiding bubbles. The cuvette was transferred to the Zetasizer’s detection slot and run. Polydispersity Index (PDI) and Z-Average (nm) were analyzed utilizing Zetasizer software and reported.

### 5.12 BVP ELISA

The method used was adopted from Hotzel 2012 [29]. Baculovirus ELISA plates were prepared with blank baculovirus (1:500 dilution), 50 *µ*L/well, and incubated overnight at 4°C. BVP dilution was made into a carbonate buffer:14.2mM Na2CO3, 34.8 mM NaHCO3, pH 9.6. BVP solution was then discarded and plates were blocked with 0.5% BSA in 1x PBS, 300 *µ*L/well for one hour at room temperature. Plates were then washed three times with 300 *µ*L per wash. All washes were performed with 1x phosphate buffered saline, pH 7.2. Antibodies were diluted to approximately 1*µ*M in 0.5% BSA in 1x PBS, and 50 *µ*L was added per corresponding well and incubated at room temperature for one hour. Plates were then washed six times with 300 *µ*L per wash. Secondary goat anti-human IgG (H+L)-HRP (Thermo 31410) secondary antibody was diluted to 0.08 *µ*g/mL in 0.5% BSA in 1x PBS and 50 *µ*L was added per well. This was incubated at room temperature for 1 hour. Plates were then washed six times, with 300 *µ*L of buffer per wash. TMB solution was prepared by mixing solution A and solution B (TMB color liquid, Sino proprietary formulation) in a 1:1 ratio, and 200 *µ*L was added per well. Color was developed for 10 minutes at room temperature. 50 *µ*L/well of 2M sulfuric acid stop solution was added, and absorbance 450 nm was immediately measured on a plate reader. BVP score is calculated via normalizing absorbance by control wells with no test antibody. A BVP score greater than 5 prognosticates a risk of faster clearance in serum of cynomolgus monkey [30].

### 5.13 Defining flag thresholds

We applied a widely used approach to define flag thresholds for each biophysical property [8, 10], which considers four property groups: self-association, hydrophobicity, polyreactivity, and stability (we consider thermostability). [8, 10] proposed thresholds by characterizing a large number of biosimilars and defining the worst 10% in each assay to be red-flagged for potential poor behavior. To translate these thresholds to our own assays, we reproduced and characterized 20 biosimilars from their study (19, after dropping one with insufficient sample purity). Our NanoDSF assay agreed with the published DSF Fab melting points for our biosimilars, therefore we used the standard definition of >64.2°C Fab as a flag cutoff. For VHHs, we utilized a VHH cutoff of >60°C *T_m_*, which was previously reported as a cutoff for low-*T_m_* VHHs [14]. Our HIC-HPLC assay proved slightly more sensitive than [8], so we scaled the retention time flag accordingly. We found the scaling factor by computing the ratio of Jain reported values versus our assay value for each of the 19 benchmark biosimilars and taking the average. In some cases there was no elution peak (Figure S3C) so the values were omitted from the calibration. Applying this scaling factor of 1.059 to the Jain threshold of <11.53 min gives <12.21 min as our green flag threshold for HIC retention time. For BVP ELISA, we took the same approach, scaling our more sensitive assay from the Jain reported 5.30 BVP score to <29.8 as our cutoff BVP score. For AC-SINS, our assay conditions resulted in lower sensitivity. In addition, due to using a 2 nm step size in the absorbance readings, our values are somewhat less granular. Because the distribution was mode-centered at 0 nm shift, ratiometric normalization (as used for HIC-HPLC and BVP ELISA) was not applicable. We therefore established a conservative AC-SINS flagging threshold of < 3.0 nm. These assay calibrations are plotted in Figure S1A, where we show the Jain flag threshold values with vertical dashed lines and the Chai-defined threshold values with horizontal dashed lines. We note that, aside from one point each in the BVP ELISA and AC-SINS assays, no other calibration points fall within the lower-right quadrants, indicating that our thresholds produce minimal false negatives.

### 5.14 *In silico* scoring

Humanness of designs was evaluated using the promb package https://github.com/MSDLLCpapers/promb following methodology proposed in BioPhi [31]. Chemical liability risk sites were identified following the Liability Antibody Profiler approach [32], with two modifications: only high-severity liabilities were included, and regions containing three consecutive hydrophobic residues were also classified as liabilities.

### 5.15 Pathhunter Agonism Assays

The GPCR Arrestin PathHunter agonism assays were performed according to standard procedures. PathHunter cell lines were expanded from freezer stocks and then seeded at 20 *µ*L per well in 384-well microplates, followed by incubation at 37°C. Intermediate dilution of sample stocks was performed to create a 5X sample in assay buffer, 5 *µ*L of which is added to the cells and incubated at 37°C or room temperature for at least 120 minutes, with the final assay vehicle concentration being 1%. Signal detection was achieved by a single addition of 15 *µ*L (50% v/v) of PathHunter Detection reagent cocktail, followed by a one-hour room temperature incubation, and then the microplates are read using a PerkinElmer Envision instrument for chemiluminescent signal. Compound activity was analyzed using CBIS data analysis suite, calculating the percentage activity as

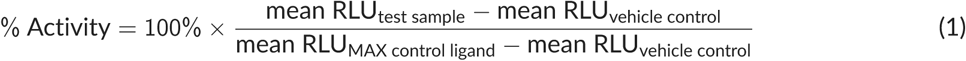

For cAMP PathHunter assays, cell expansion and seeding was carried out in the same way. cAMP modulation was determined using the DiscoverX HitHunter cAMP XS+ assay. For agonist determination, cells were incubated with sample in the presence of EC80 forskolin to induce response. For G_i_ cAMP assays, the following forskolin concentrations were used: CXCR4 cAMP: 20 *µ*M Forskolin, CXCR6 cAMP: 15 *µ*M Forskolin. Media was aspirated from the cells and replaced with 10 *µ*L HBSS/10mM Hepes. An intermediate dilution of sample stocks was performed to generate 4X sample in assay buffer, and 5 *µ*L of this 4X sample was added to the cells. Subsequently, 5 *µ*L of 4X EC80 forskolin diluted in HBSS/HEPES was added, and the plates were incubated at 37°C for 30 minutes. The final assay vehicle concentration was 1%. After appropriate compound incubation, assay signal was generated through incubation with 5*µ*L cAMP XS+ Ab reagent followed by 20 *µ*L cAMP XS+ ED/CL lysis cocktail for one hour at room temperature. 20 *µ*L cAMP XS+ EA reagent was added for two hours at room temperature. Microplates were read following signal generation with a PerkinElmer Envision instrument for chemiluminescent signal detection. Compound activity was analyzed using CBIS data analysis suite (ChemInnovation, CA). CXCR4 and CXCR6 activity are measured via G_i_-coupled cAMP level reduction, calculating the percentage activity as

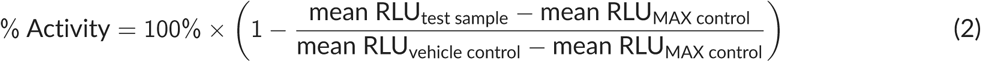

## A Supplementary information

**Figure S1.**
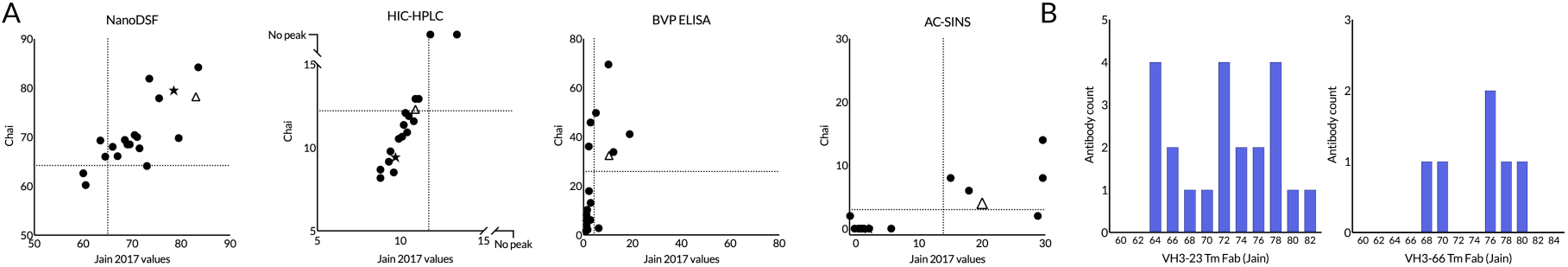
Calibration of developability assays on control antibodies. (A) Correlation between values reported in the Jain 2017 dataset [8] and values measured by Chai on the same set of control antibodies, for four developability criteria. Aside from one point each in the BVP ELISA and AC-SINS assays, no calibration points fall within the lower-right quadrants, indicating that our thresholds introduce minimal false negatives. (B) Reported Fab melting points for molecules with VH3-23 and VH3-66 frameworks in the Jain 2017 dataset [8].

**Figure S2.**
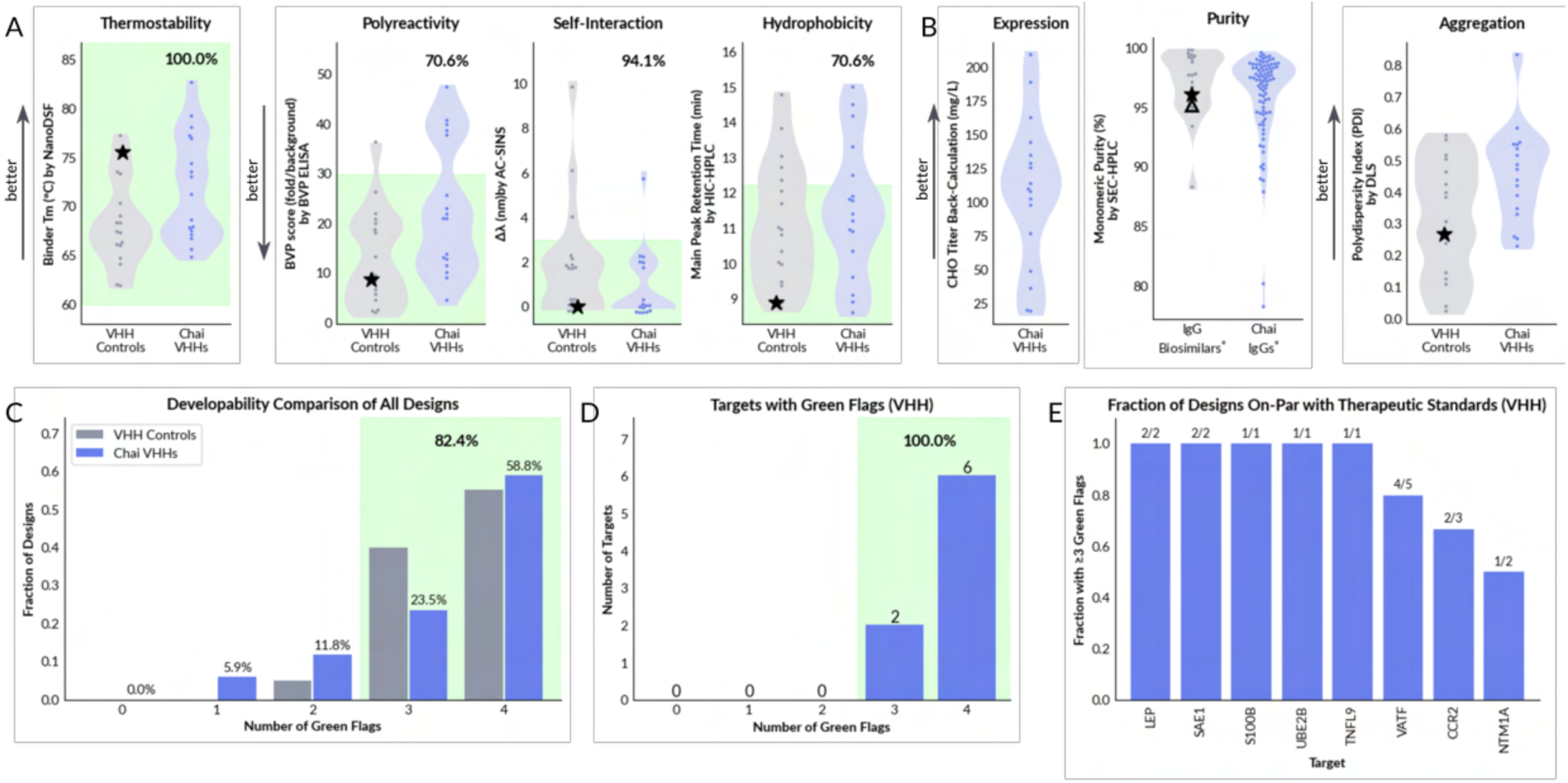
Developability assay panel for VHH designs. (A) Core developability properties measured for VHH designs. (B) Additional developability lab measurements for VHH designs. (C) Number of green flags as fraction of overall designs and biosimilars. (D) Number of green flags of best design per target. (E) Fraction of designs that have 3 or more developability green flags by target, annotated with absolute numbers. Caplacizumab (star) serves as an example of a VHH with a strong developability profile.

**Figure S3.**
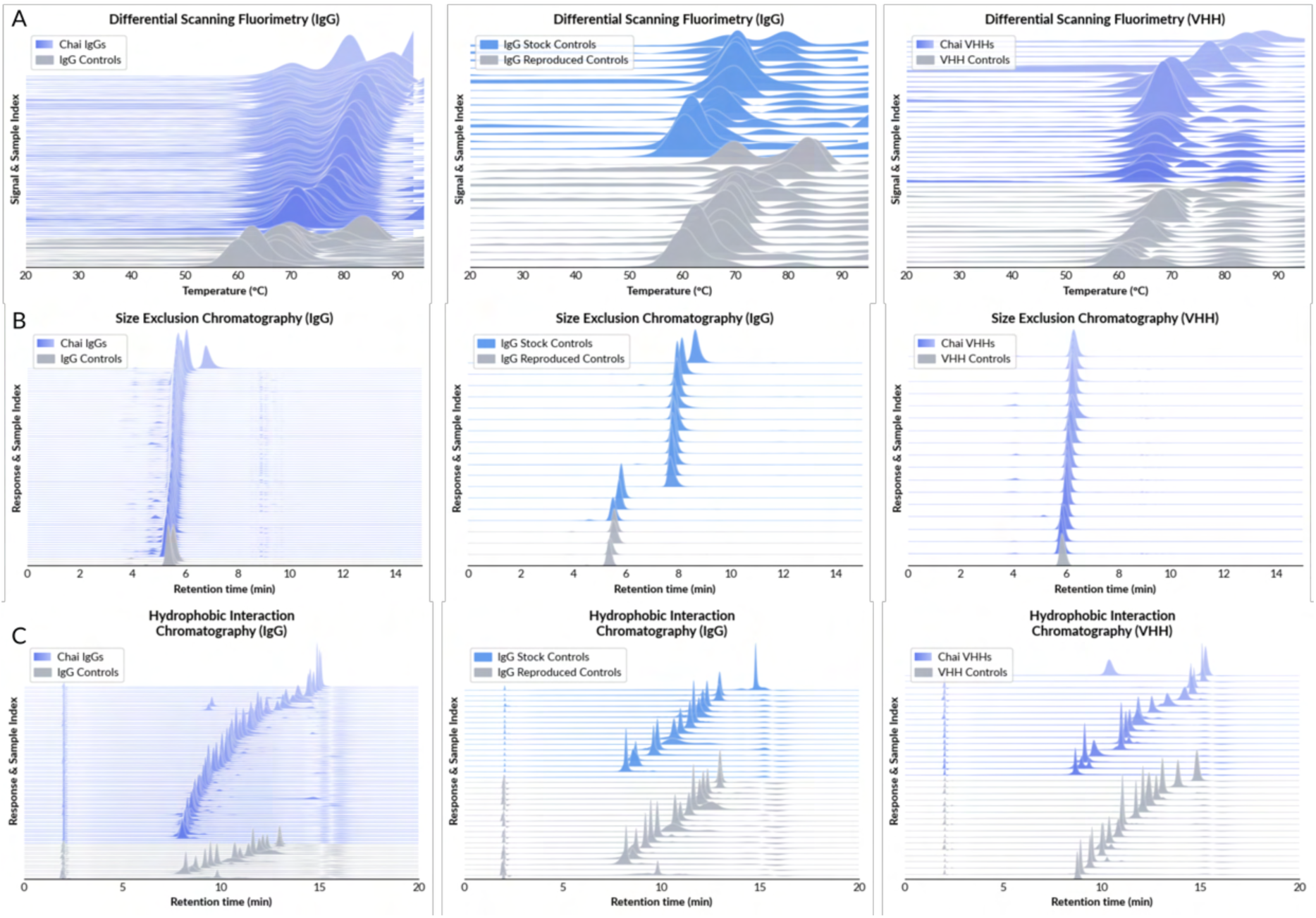
Ridge plots for raw data of Chai IgGs, reproduced controls, IgG stock controls and Chai VHHs in DSF (A), SEC-HPLC (B), and HIC-HPLC (C) assays. “IgG Controls” are the same as the IgG reproduced controls. IgG stock controls are data collected on highly-polished biosimilar protein lots. Only samples passing a 90% purity cutoff are plotted.

**Figure S4.**
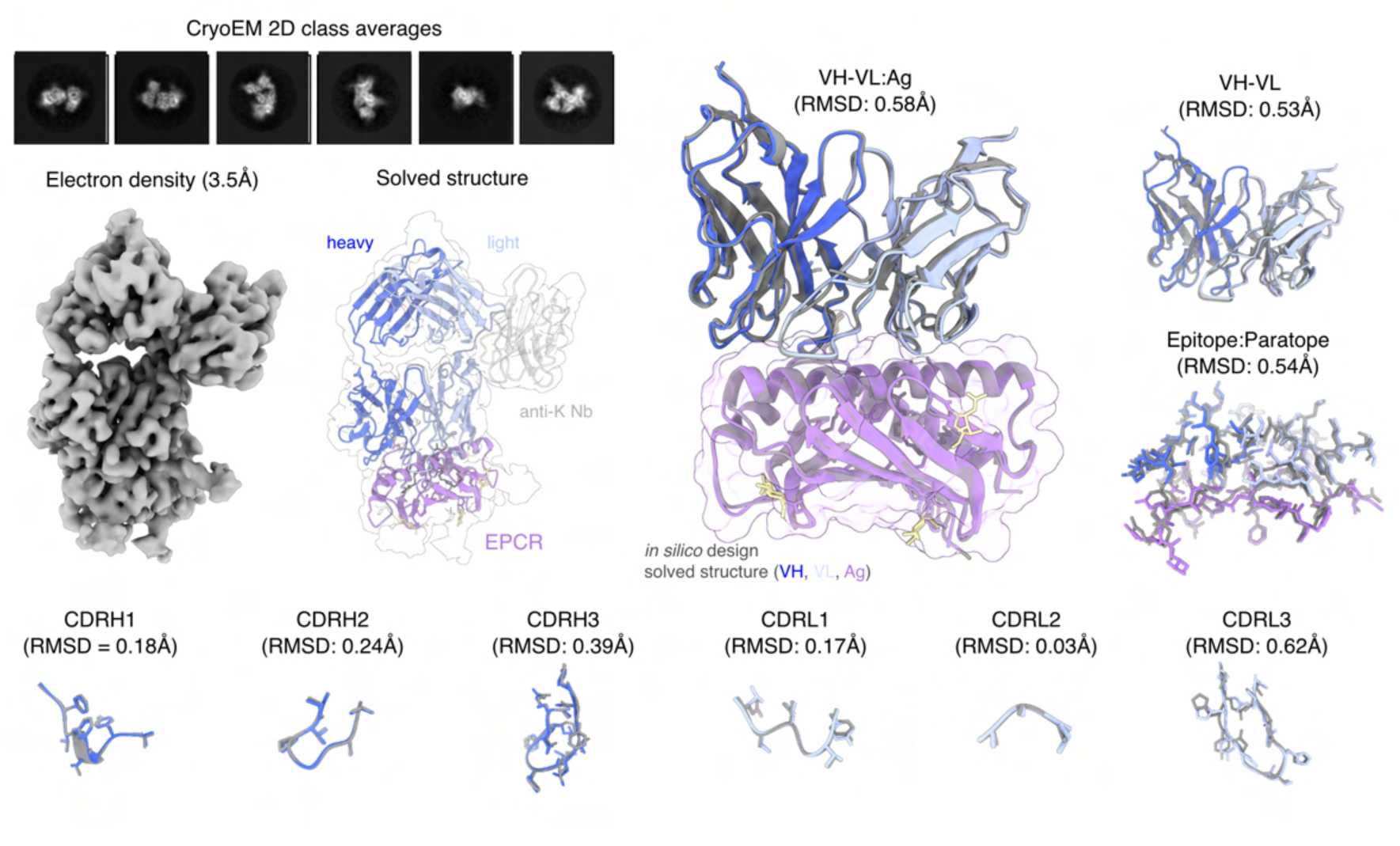
Extended cryo-EM data for design against EPCR. Predicted structures are shown as cartoons in grayscale while experimentally resolved are colored.

**Figure S5.**
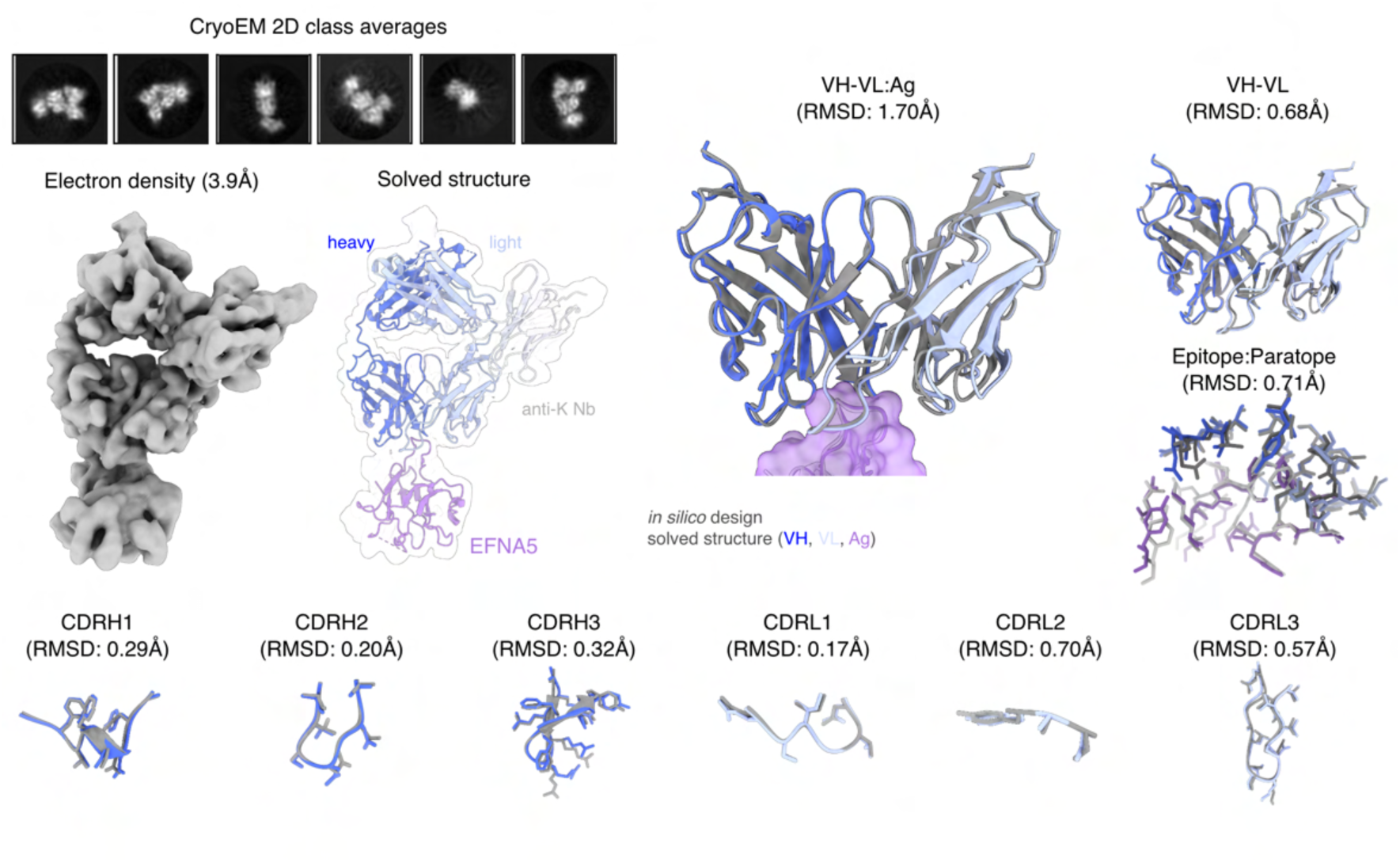
Extended cryo-EM data for design against EFNA5. Predicted structures are shown as cartoons in grayscale while experimentally resolved are colored.

**Figure S6.**
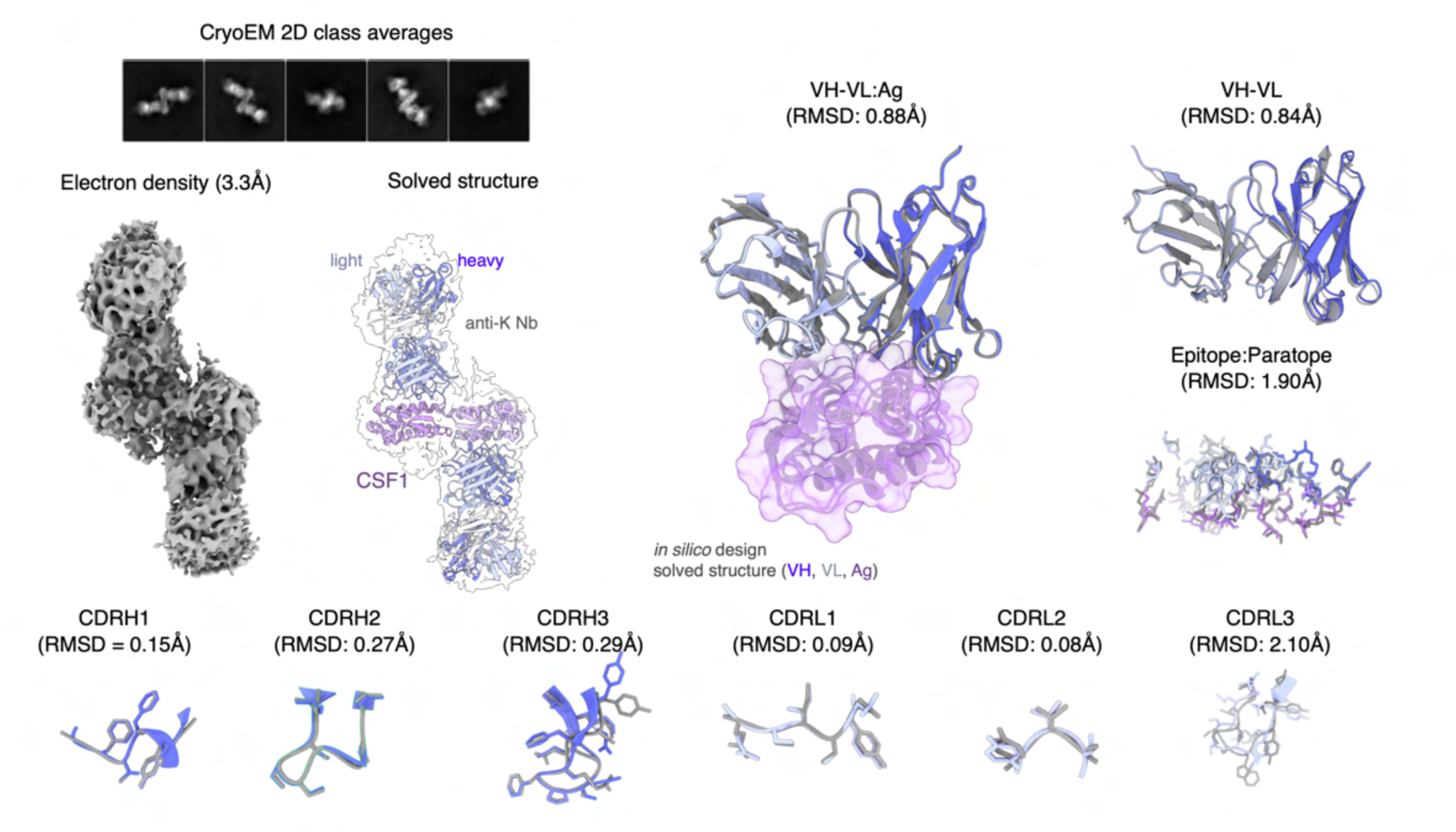
Extended cryo-EM data for design against CSF1. Predicted structures are shown as cartoons in grayscale while experimentally resolved are colored.

**Figure S7.**
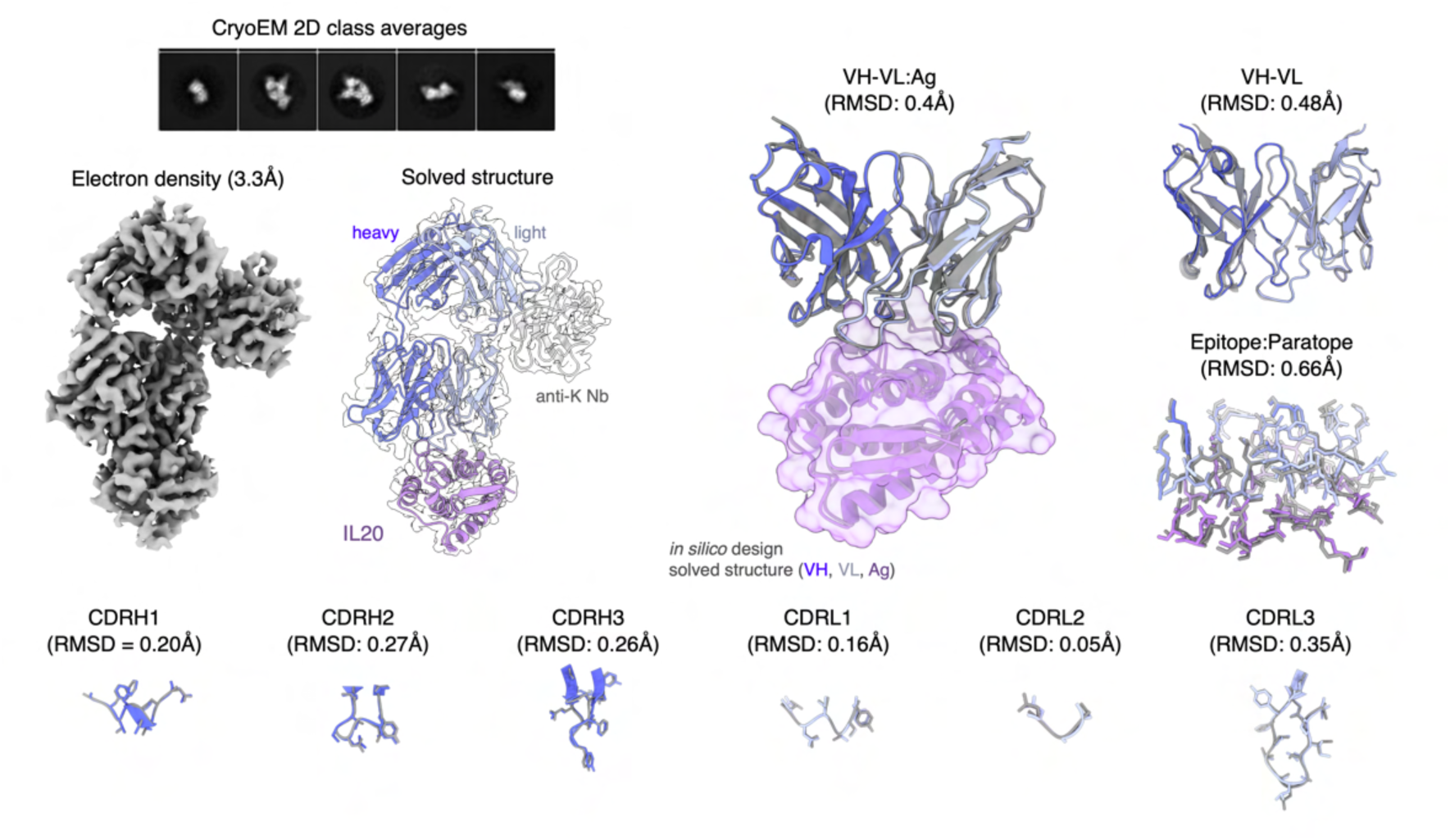
Extended cryo-EM data for design against IL20. Predicted structures are shown as cartoons in grayscale while experimentally resolved are colored.

**Figure S8.**
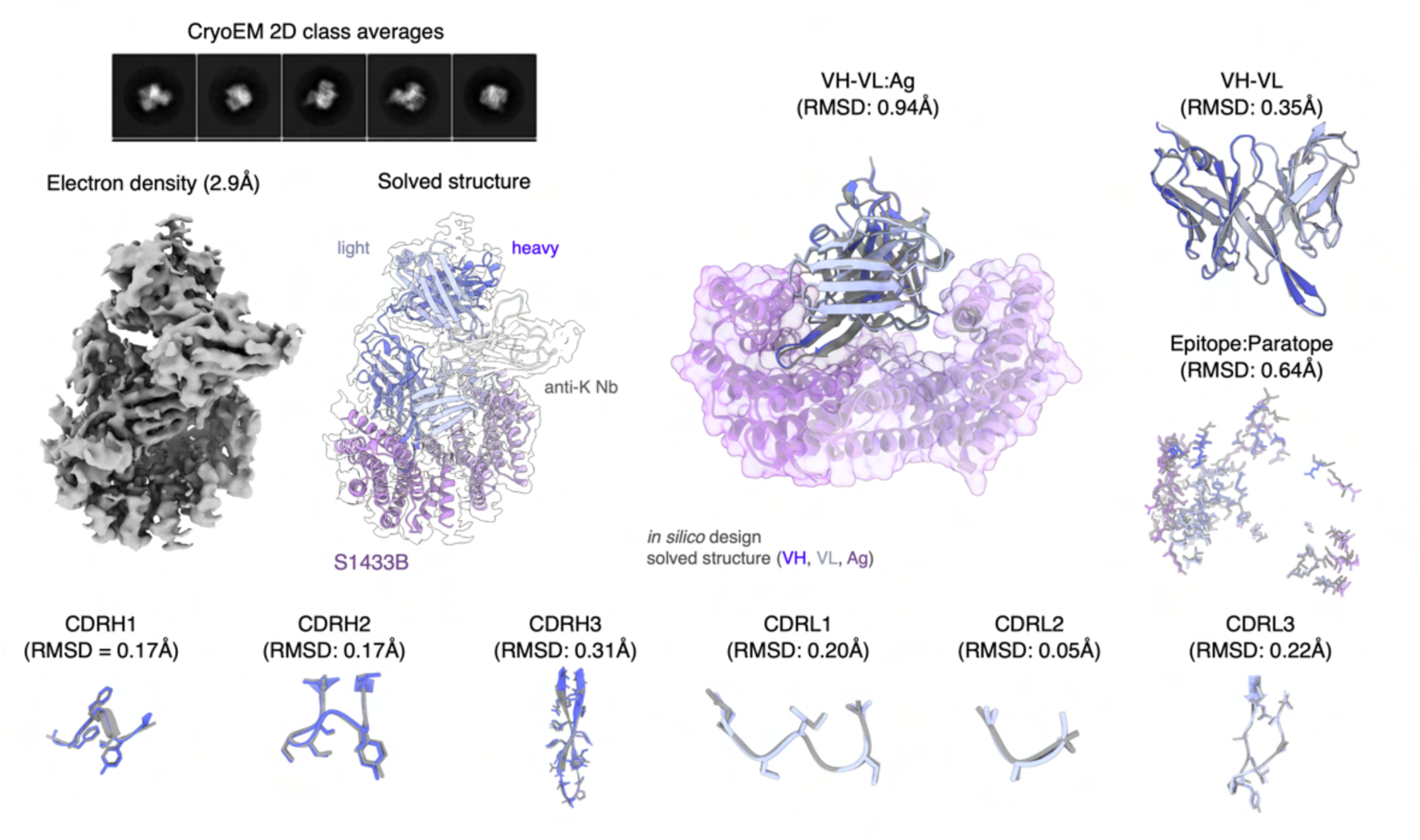
Extended cryo-EM data for design against S1433B. Predicted structures are shown as cartoons in grayscale while experimentally resolved are colored.

**Figure S9.**
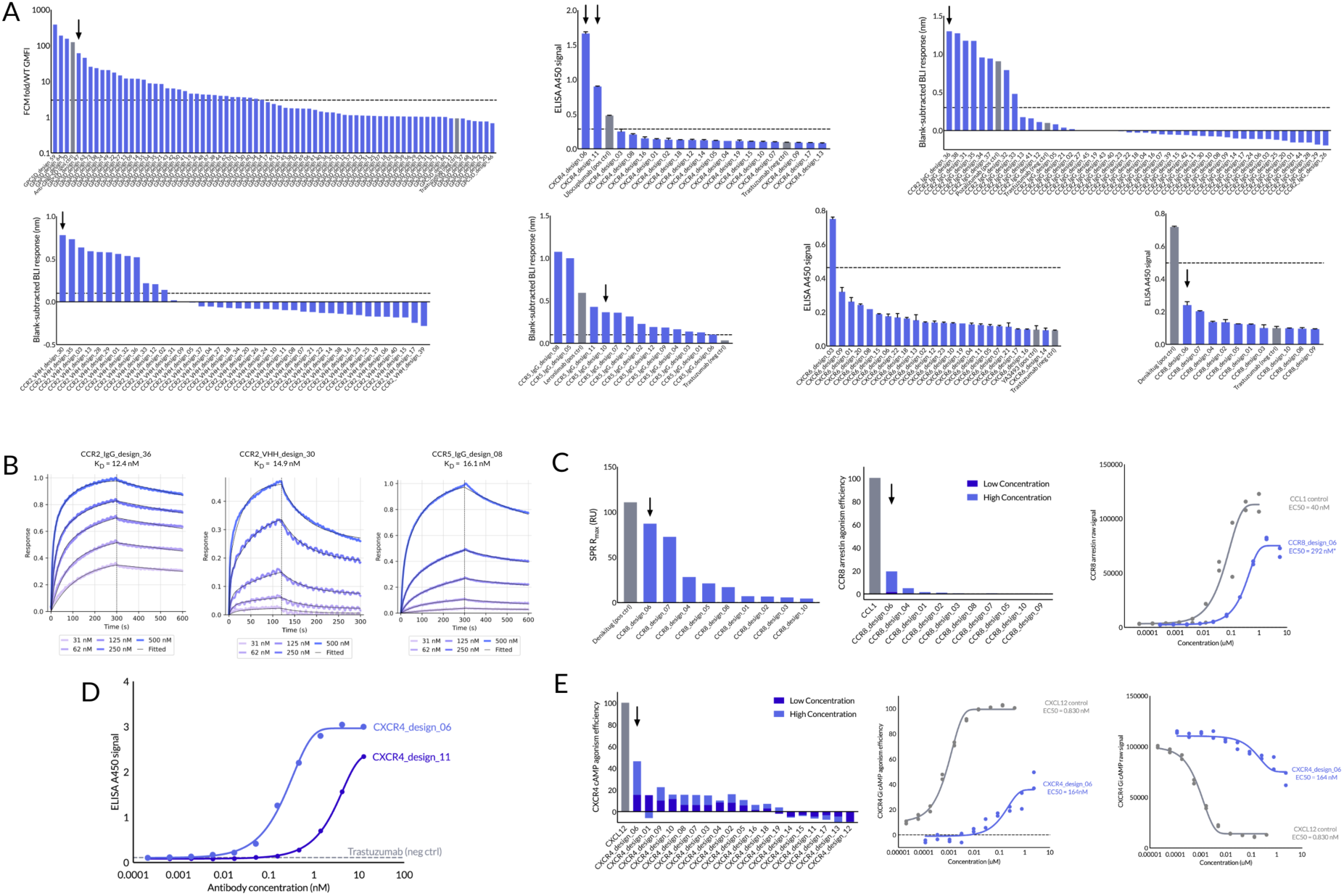
Binding and functional characterization of GPCR binders. (A) Primary hit screening was performed as described (Table 2). Flow cytometry screening was performed across 73 GPRC5D designs yielding 35 hits. Arrow indicates GPRC5D_design_47 featured in Figure 4A-C. VLP ELISA screening of 19 CXCR4 designs was performed, yielding 2 hits. Arrows indicate the two clones selected for follow-up multipoint ELISA screening (Panel D) VLP ELISA screening was performed across 23 CXCR6 designs, yielding 1 hit. BLI screening was performed across 40 CCR2 IgG and 45 VHH designs, yielding 10 and 8 hits respectively. Arrows indicate the clones selected for monovalent BLI kinetics plotting (Panel B). BLI screening of CCR5 IgG designs was performed, yielding 13/13 positive hits. Arrow indicates the clone selected for monovalent BLI kinetics plotting (Panel B). VLP ELISA screening was performed across 10 CCR8 designs, yielding no hits in this assay. (B) Monovalent affinities were determined for representative designs as indicated via arrows in FigureS9A. CCR2_-IgG_design_36’s affinity was determined to be 12.4 nM, and CCR2_VHH_design_30’s affinity was determined to be 14.9 nM via binding to nanodisc-solubilized CCR2. CCR5_IgG_design_08’s affinity was determined to be 16.1 nM via binding to nanodisc-solubilized CCR5. (C) CCR8 VLP screening was followed up by SPR with nanodisc-solubilized CCR8 protein. Two-point PathHunter CCR8 arrestin agonism screening was performed across 9 designs with CCL1. CCR8_design_06 was subsequently evaluated via full dose range titration in the same PathHunter assay. Raw CCR8 arrestin luminescence values were plotted which form the basis of the percent agonism efficiency plot in Figure 4F. (D) Two CXCR4 VLP ELISA hits identified in Panel A, CXCR4_design_06 and CXCR4_design_11, were subsequently analyzed via multipoint VLP ELISA screening. (E) All CXCR4 clones were assayed via two-point PathHunter CXCR4 cAMP agonism screening. Following this, CXCR4_design_06 was then assayed in a full dose range titration compared to CXCL12 ligand. Both percent agonism and raw luminescence values were plotted, to which a 4-parameter nonlinear fit was performed to determine the EC_50_ values.

**Figure S10.**
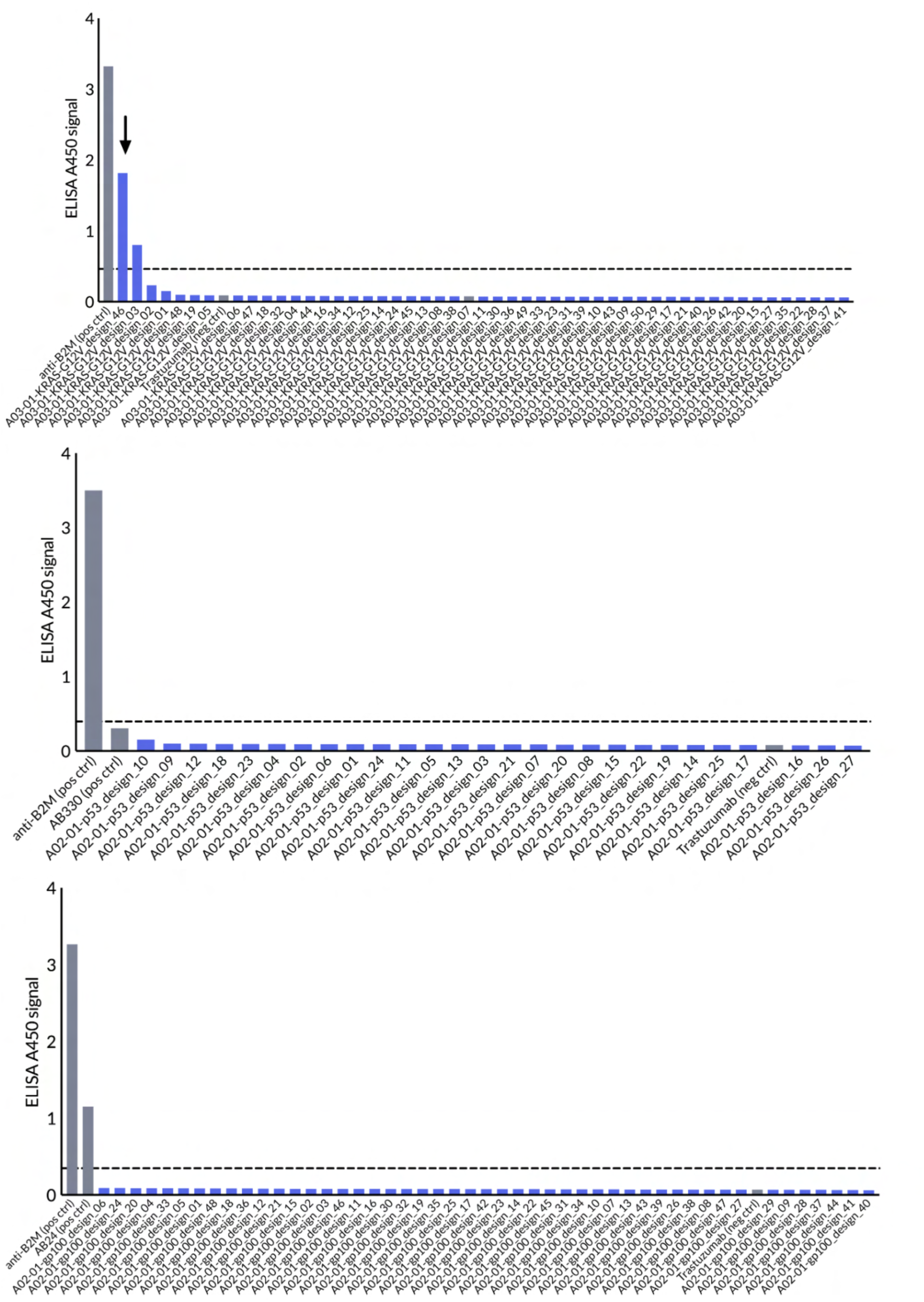
Binding screening of pMHC binders. ELISA screening was performed across 50 HLA-A*03:01 KRAS-G12V designs. Design-46 depicted in Figure 5A is indicated with an arrow. ELISA screening was performed across 27 HLA-A*02:01 p53-R175H designs. The AB330 positive control failed to demonstrate appreciable binding. ELISA screening was performed across 48 HLA-A*02:01 gp100 designs.

**Figure S11.**
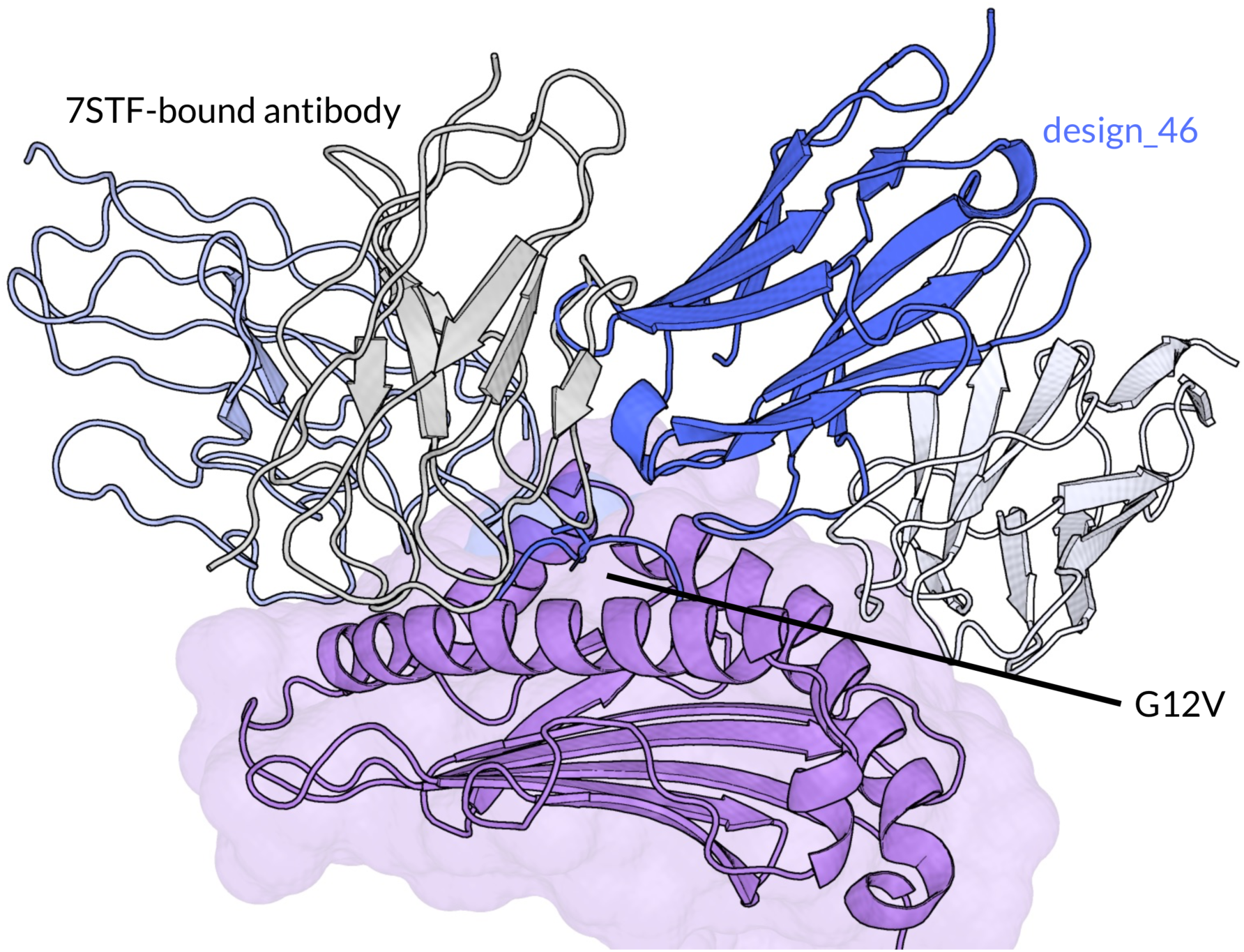
Structure modeling of pMHC binder using Chai-2. Structural comparison of HLA-A*03:01 KRAS-G12V design-46 predicted binding mode compared to antibody fragment V2 bound to PDB: 7STF. Design_46’s VH is depicted in dark blue and VL in white, and V2’s VH is depicted in light blue and VL in gray. The predicted binding mode of the Chai designs is different from the 7STF-bound antibody.

### A.1 Tables of reagents

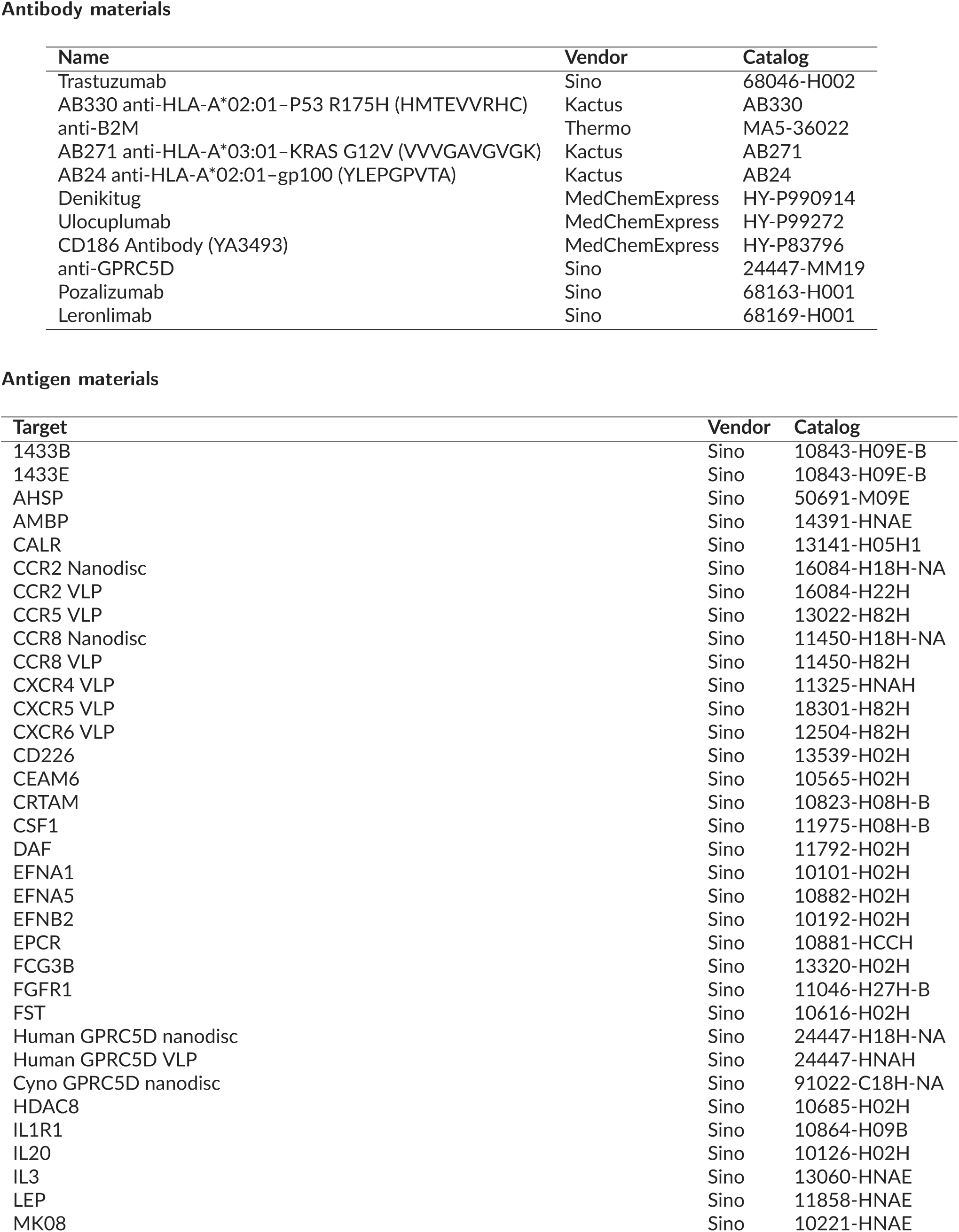
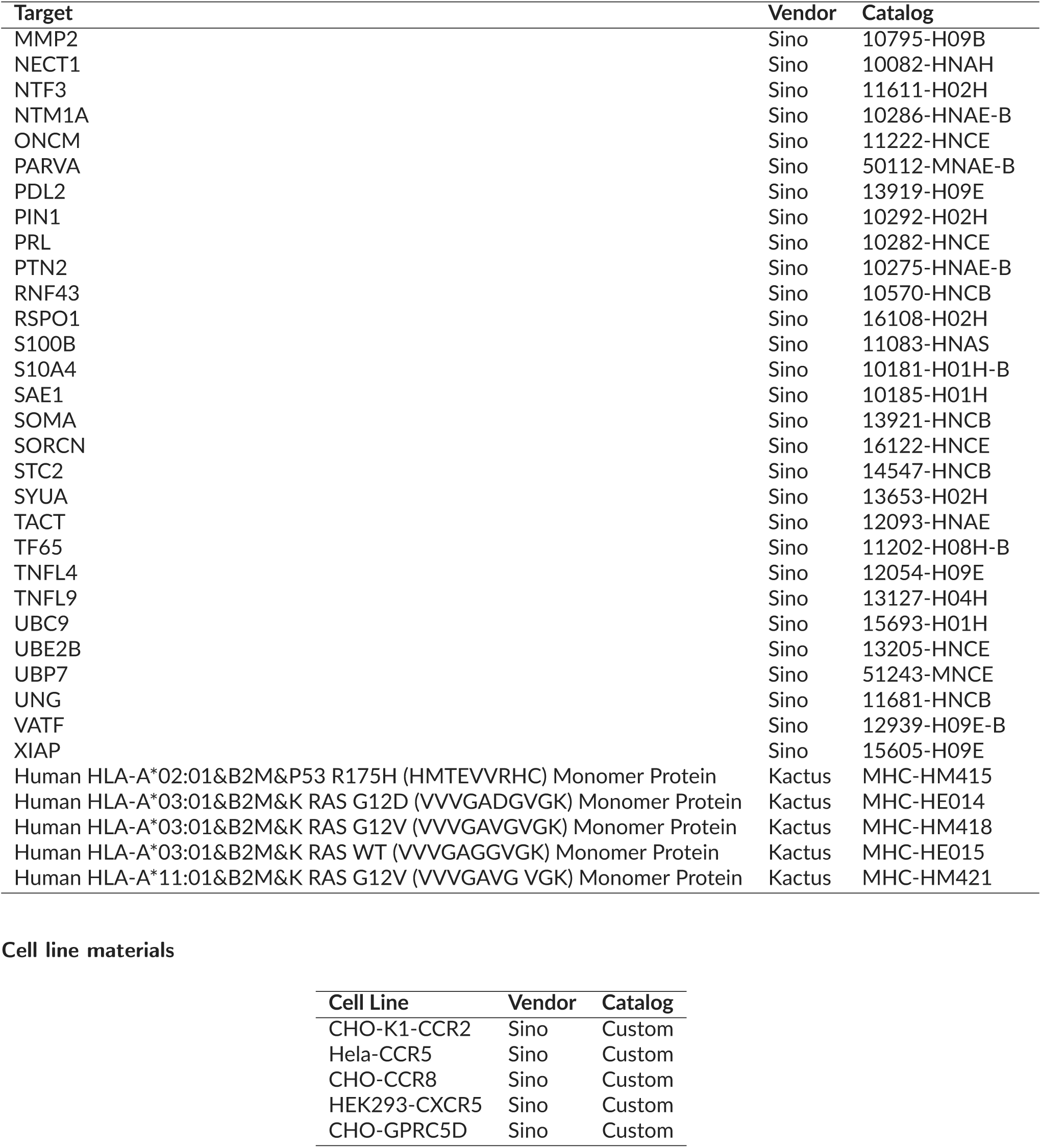

